# Neural bases of space-specific distractor biases in visual working memory

**DOI:** 10.1101/2023.12.24.573161

**Authors:** Deepak V Raya, Sanchit Gupta, Devarajan Sridharan

## Abstract

Information held in working memory (WM) is remarkably resilient to distraction. Yet, perceptual distractors that share mnemonic features can impact WM profoundly; the neural basis of this phenomenon remains unclear. With multivariate decoding of human electroencephalography recordings, we investigate how delay-period perceptual distractors bias WM. Participants memorized the orientations of cued and uncued grating memoranda that appeared in opposite hemifields. A grating distractor, flashed during the delay period, produces space-specific biases: memorized features are attracted towards or repelled away from the distractor’s orientation depending, respectively, on when the distractor appeared in the same hemifield as the memorandum, or opposite to it. Neural prioritization in WM by cueing, and stronger memorandum maintenance mitigate this bias, whereas stronger distractor encoding enhances it. Lastly, a ring-attractor model with cross-hemifield inhibition mechanistically explains the origins of these spatially-antagonistic biases. Our results reveal how lateralized sensory buffers critically enable perceptual distractors to bias visual WM.

**Lay Summary:** Working memory (WM) – the ability to momentarily store important items – is remarkably robust to distraction. Yet, when a salient object with features resembling the memorized item (“perceptual distractor”) appears in the environment, it can alter WM appreciably. What neural mechanisms render WM resilient to distraction, and how can perceptual distractors affect it so profoundly? We address this question by presenting a salient distractor, at unpredictable times, during the WM delay-period. Surprisingly, the distractor’s bias on WM depends on its proximity to the memorandum’s original location. With state-of-the-art neural decoding and computational modeling, we identify mechanisms that mediate these space-specific distractor effects. The findings advance our understanding of WM’s resilience to distraction and may inform cognitive therapies for treating WM deficits.

## Introduction

Working memory (WM) is a remarkable cognitive ability that enables us to transiently hold and manipulate information that is no longer present in the environment. Mnemonic information in WM must be maintained in the absence of external sensory input, and it is increasingly evident that early sensory areas are actively involved in such maintenance^1–5^. As a result, mnemonic information may be susceptible to interference by new sensory inputs, such as salient “perceptual” distractors^5–7^. Yet, a significant body of literature suggests that information in WM is surprisingly resilient to distraction^5,6,8,9^. For example, behavioral performance in a WM task involving oriented gratings was robust, despite multiple task-irrelevant (face or place) distractors being presented during the delay period^6^. Similar results have been observed in many recent studies of visual WM^5,10^.

Despite this robustness, recent evidence suggests that perceptual distractors may interfere systematically, with the content of WM^5,8,10,11^. In particular, some distractors are more potent than others at “biasing” the content of working memory, especially during the delay-period ^8,9,11–16^. The extent to which a distractor’s features match task-relevant features of the memorandum determines the strength of its bias, with greater feature similarity yielding greater bias in WM^8,15^. Typically, such biases are “attractive” in that the memorandum’s report is biased towards the distractor’s features^8,10,11,17^. For instance, viewing a task-irrelevant orientation during the WM delay period biases the remembered target’s orientation towards the distractor’s orientation^10,17^. Similarly, spatial distractors can bias the remembered location towards the distractor’s location^11^. Such perceptual distractor biases have been reported even for complex features. For example, remembered face representations are more strongly biased by the presentation of face image distractors than by scene image distractors^15^.

Identifying mechanisms that mediate distractor interference, including perceptual biases, in WM representations is a topic of active research^5,8–11^. Previous studies measured brain responses during the WM delay epoch, and have found evidence for a distributed representation of mnemonic information in diverse brain regions^18^, including sensory (visual) cortex^1–5^, posterior parietal cortex (PPC, ^10,19,20^), the intraparietal sulcus (IPS, ^6,10,21^), as well as in the prefrontal cortex (PFC, ^7,21,22^). Converging evidence from these studies suggests that while mnemonic information in sensory cortex may be susceptible to distractor interference and bias, information in higher cortical areas, including PPC^6,10^ and PFC^9,23^, is much more robust to such interference. For example, when a distractor grating with a random orientation was presented during the delay period of a working memory task, it produced biases in neural representations in the early visual cortex (V1-V3), but not in the IPS^10^. A related study showed that the distractor’s orientation affected neural representations in early visual cortex (V1-V4), but higher cortical regions, including the IPS and lateral occipital cortex, were unaffected^5^. In fact, such stability of representations in higher cortical areas is hypothesized to mediate the remarkable resilience of WM to sensory distractors ^8,24^.

Despite this emerging literature, a key question remains unanswered: what neural mechanisms govern how a perceptual distractor affects the contents of working memory during maintenance (“delay-period perceptual distraction”^25^)? Previous studies on distractor effects overwhelmingly employed simple WM task designs: for example, most tasks involved a single, central distractor presented at fixed times during the delay epoch^5,8–12,14–16,26^. Moreover, many more previous studies investigating mnemonic bias employed functional magnetic resonance imaging (fMRI)^5,10,11^ to study WM biases in the brain. The poor temporal resolution of fMRI precludes addressing fundamental questions about rapid, distractor timing effects: Because the memorandum and distractor occur within a few hundred milliseconds of each other, their evoked blood oxygenation level dependent (BOLD) responses evolve over the timescale of several seconds and are challenging to disentangle. No previous study, to our knowledge, has explored distractor-induced bias effects by varying the distractor’s location and its timing relative to the memorandum, systematically.

Here, we addressed this question with multivariate stimulus decoding of high-density human electroencephalography (EEG) recordings, which enabled measuring distractor’s neural effects with millisecond precision. We characterized how perceptual distractors induce mnemonic biases in participants (n=24) performing a continuous response, visual WM task involving memorandum orientation recall (Fig. 1A). We show that a task-irrelevant distractor grating presented during the delay period produces either an attractive or a repulsive bias in features stored in WM depending on the distractor’s proximity to the memorandum’s location. With state-of-the-art neural decoding, we show that robust maintenance and input gating strongly shield mnemonic information from distractor interference. Lastly, we explain the mechanistic origin of these biases with a two-tier ring attractor model^27,28^.

**Figure 1.**
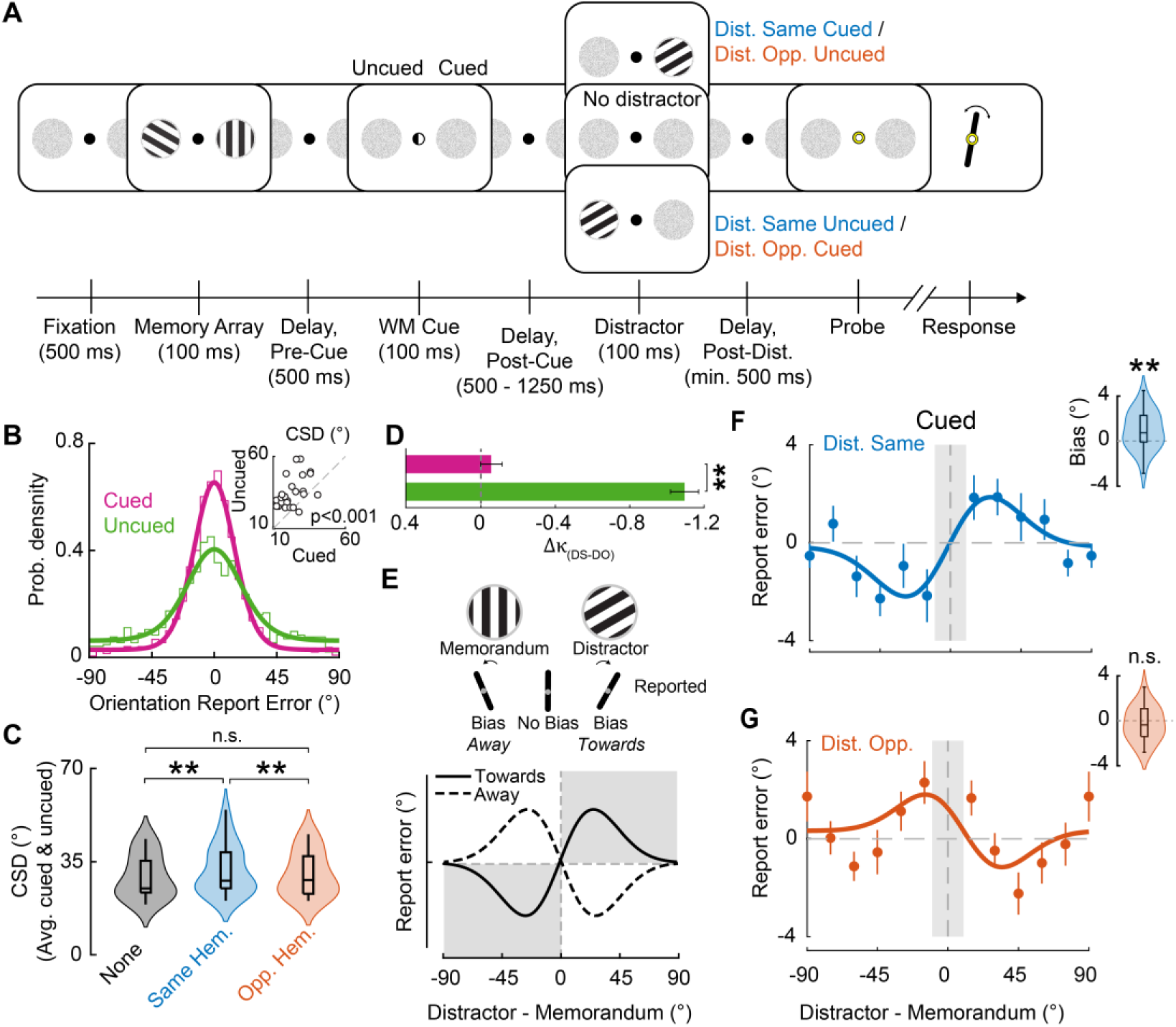
Perceptual distractor produces antagonistic biases during recall. **A.** Visual working memory task with a distractor. Two randomly oriented gratings were flashed briefly, one in each hemifield. A retro-cue (white semicircle) indicated the location of the erstwhile grating that would likely be probed for response (70% validity). Following a variable delay-period, on 80% of the trials, a singleton perceptual distractor grating was flashed either on the right or left hemifield (upper and lower rows); on the remaining trials (interleaved) no distractor appeared (middle row). At the end of the trial, a central response probe (white or black) indicated the location of the grating (cued or uncued, respectively) to be recalled and reported. **B.** Distribution (thin histogram) of the orientation report errors for the cued (magenta) and uncued (green) trials (n=24, data pooled across participants). Thick lines: von Mises fits. (*Inset*) Circular standard deviation (CSD) of the report errors for “no distractor” trials, for cued (x-axis) versus uncued (y-axis) grating reports. Dashed line: line of equality. Data points: individual participants. **C.** CSD for no distractor trials (black), when the distractor appeared in the same (blue), or opposite (orange), hemifield as the probed memorandum. Violin plots: kernel density estimates; center line: median; box limits: upper and lower quartiles; whiskers: 1.5x interquartile range. **D.** Change in precision between the distractor-same and distractor-opposite trials (Dk=k_DistSame_– k_DistOpp_) for the cued (magenta) and uncued (green) trials. Error bars: Jackknife. **E.** Schematic of bias. (*top*) The distractor (left grating) may bias the memorandum’s (right grating) recalled orientation away from its own (left bar), towards its own (right bar), or not at all (middle bar). (*bottom*) Error in the probed memorandum’s orientation (signed) (y-axis) as a function of its orientation relative to the distractor (x-axis). Positive and negative values (both axes): clockwise and counter-clockwise relative orientations, respectively. Solid and dashed curves: attractive bias (toward), or repulsive bias (away from), the distractor’s orientation, respectively. **F.** Average behavioral bias (n=24) when the distractor appeared in the same hemifield as the cued memorandum. Solid line: Derivative of Gaussian fit (Methods). Data smoothed over three successive orientation bins for visualization purposes only. Error bars: s.e.m. Gray shaded rectangle: untested relative orientation values (±10°). (*Inset, top right*) Violin plot: bias strengths across participants; positive (negative) values denote attractive (repulsive) bias. Other conventions are the same as in panel C. **G.** Same as in panel F, but for trials in which the distractor appeared in the hemifield opposite to the cued memorandum. Other conventions are the same as in panel F. (All panels) *: p<0.05; **: p<0.01; n.s.: not significant.

The results show how lateralized sensory buffers could provide a conduit by which task-irrelevant sensory input that is perceptually similar to remembered items could gain access to WM and bias its mnemonic content. Our study also provides a general computational framework to study the resilience of visual working memory to perceptual distractor effects.

## Results

### Distractors induce space-specific biases in visual working memory

We tested n=24 human participants on a visual WM task involving grating orientation report (Fig. 1A, Methods). Following fixation, two randomly oriented gratings (“memory array”) were flashed briefly (100 ms) one in each visual hemifield. A brief interval (500 ms) after grating offset, a spatial retro-cue (filled white semi-circle) appeared, which indicated the hemifield of the erstwhile grating that would likely be probed for response (cue validity: 70%). A fixed epoch (2450 ms) after the memory array offset, a probe appeared (central filled circle) whose color (white or black) indicated the grating (cued or uncued, respectively) whose orientation the participants must reproduce (Methods).

To study the effect of perceptual distractors in visual WM, a singleton, distractor grating – randomly oriented but otherwise identical to the stimuli in the memory array (Methods) – was flashed briefly (100 ms) at a random time (500-1250 ms, exponentially distributed) during the delay period after retro-cue offset. The distractor was presented in 80% of the trials – interleaved pseudorandomly with 20% “no distractor” trials – and appeared either on the left or the right hemifield with equal probability. We refer to the retro-cued hemifield, the grating at this location or its representation in WM as the “cued” hemifield, grating, or representation, respectively; their counterparts in the opposite hemifield are referred to as the “uncued” hemifield, grating or representation, respectively.

First, we tested if participants were indeed utilizing the retro-cue for selection in visual WM, including only trials in which no distractor appeared (20% of all trials). The error in orientation reports, as measured by the circular standard deviation (CSD) of the orientation report error, was significantly lower on cued trials compared to uncued trials (CSD_Cued_=23.4°±1.6° [mean±s.e.m.]; CSD_Uncued_=35.2°±2.2°, p<0.001, Wilcoxon signed rank test, Bayes Factor (BF_10_) >10^2^; n=24 participants) (Fig. 1B), confirming robust selection of the retro-cued representation in visual WM.

Next, we analyzed the effect of the perceptual distractor on the cued and uncued gratings’ orientation reports including all trials (80%) in which the distractor was presented. We tested for spatially lateralized effects; specifically, we hypothesized that the precision of orientation reports would be different depending on whether the distractor was presented in the same (“distractor-same” condition) or on the opposite hemifield (“distractor-opposite” condition) as the probed stimulus. We found robust evidence confirming this hypothesis: a 2-way ANOVA with the CSD of orientation errors as the dependent variable revealed a significant main effect of distractor location (F_(1,23)_=11.06, p<0.001), but no significant interaction effect between cueing and distractor location (F_(1,23)_=0.48, p=0.625). Post-hoc analyses revealed a significant increase in error for distractor-same compared to no-distractor trials (Fig. 1C, CSD_NoDist_=29.3°±1.7°, CSD_DistSame_=32.0°±1.9°, p=0.002, BF=40.34, signed rank), but not for distractor-opposite compared to no-distractor trials (Fig. 1C, CSD_NoDist_=29.3°±1.7°, CSD_DistOpp_=30.3°±1.7°, p=0.103, BF=1.16, signed rank); data pooled across cued and uncued locations).

Lastly, we analyzed the degradation in the precision of orientation reports across the cued and the uncued locations, induced by the distractor. For this, we pooled the orientation reports across all participants and fit a von Mises and uniform mixture model^29^ to behavioral reports, separately for distractor-same and distractor-opposite trials; the degradation in the accuracy of orientation reports induced by the distractor was quantified with the difference in precision (Δκ) across these two conditions (κ_DistSame_–κ_DistOpp_) (Methods). Δκ was significantly greater on the uncued hemifield (Δκ_uncued_= −1.09±0.075 than the cued hemifield (Δκ_cued_=-0.06±0.057, p=0.002, permutation test) (Fig. 1D). Essentially identical results were obtained using the modulation index of the precision, quantified as MI-κ = (κ_DistSame_–κ_DistOpp_)/ (κ_DistSame_+κ_DistOpp_) (p=0.003, permutation test).

In other words, the distractor affected the precision of orientation reports for the memorandum when it occurred, subsequently, in the same hemifield, and this effect was stronger for the uncued memorandum, as compared to the cued memorandum.

It is possible that the increase in orientation report error was merely a consequence of a generic disruption of WM maintenance by the perceptual distractor – a salient, flashed stimulus. Alternatively, it is possible that the increase in error arose from a specific bias in WM during the delay induced by the distractor’s orientation. To distinguish between these possibilities, we tested whether the behavioral orientation report of the probed memorandum was biased by the distractor’s orientation, by plotting the signed orientation report error for the probed grating as a function of the difference between the distractor and the probed gratings’ orientation; bias was quantified with the signed area-under-the-curve metric^28^ (Fig. 1E) (Methods).

Orientation reports for the cued memorandum were systematically biased towards the perceptual distractor’s orientation (“attractive” bias) when the latter appeared in the same hemifield as the former (Fig. 1F) (Cued: Bias_DistSame_=0.94°±0.35°, p=0.004, BF_+0_=7.50). By contrast, when the distractor appeared in the opposite hemifield, orientation reports revealed a trend of being biased away (“repulsive” bias) from the distractor’s orientation (Fig. 1G) (Cued: Bias_DistOpp_=-0.3°±0.3°, p=0.177, BF_-0_=0.53). Although the repulsive bias effect was not statistically significant when data from all trials were pooled, in subsequent sections we show that when subsets of trials are selected based on objective criteria (e.g., distractor encoding strength) the pattern of repulsive bias becomes palpable, and significant (see next; e.g., Fig. 3 C-D).

A similar trend in attractive bias was apparent for the uncued grating also when the distractor appeared on the same hemifield as the memorandum (SI Fig. S1). Yet, both bias trends on the uncued hemifield were not statistically significantly different from zero (Uncued: Bias_DistSame_=1.76°±1.15°, p=0.067, BF_+0_=1.12; Bias_DistOpp_=0.98°±1.07°, p=0.818, BF_-0_=0.115), possibly because of the comparatively fewer trials (30%) in which uncued gratings were probed. Given the substantially greater proportion of trials probed on the cued hemifield that led to robustly measurable bias effects, we limit subsequent analyses of distractor-induced behavioral biases to cued-hemifield probed trials.

To test for the possibility that the behavioral bias arose from the observers mis-reporting the distractor’s orientation as that of the probed memorandum’s (target) on a subset of trials, we fit a model (von Mises) comprising a mixture of target and distractor orientations, as well as uniform noise, to their responses (“target+distractor” model, Methods) (SI Fig. S2A). We then compared this model against a more parsimonious model that did not incorporate the distractor orientations in the mixture (“target alone” model) (SI Fig. S2B). Models were fit for each participant individually, and formal model comparison was performed with the corrected Akaike information criterion (AICc) and Bayesian Information Criterion (BIC) that balance model complexity with goodness-of-fit^31,32^; a lower value of each metric indicates the preferred (selected) model.

AICc was significantly higher for the “target+distractor” model than for the “target alone” model (AICc: target + distractor =764.98±34.78, mean±s.e.m. across participants, target alone = 760.91±34.94, p<0.001, Wilcoxon signed-rank test) (SI Fig. S2C). These results were exactly replicated also with the BIC metric (BIC: target + distractor = 789.48±34.785, target alone = 773.26.66±34.94, p<0.001) (SI Fig. S2C). Moreover, examining the mixture proportions revealed that the mixture probability associated with the distractor component was substantially and significantly smaller (∼1/12*th*, p<0.001) than the target component in the “target + distractor” model (SI Fig. S2D). We repeated these analyses with a mixture model that did not include the random component in either the “target + distractor” or “target alone” models (von Mises alone, see Methods) and observed an essentially identical pattern of results.

In sum, a perceptual distractor presented during the delay period systematically disrupted working memory, with cued memoranda being better shielded from interference as compared to uncued memoranda. Moreover, the distractor induced antagonistic biases – attractive and repulsive – when it appeared in the same versus opposite hemifields, respectively, as the cued memorandum, during the delay-period. This space-specific pattern of biases suggests that maintenance in working memory is mediated, at least in part, by a spatially lateralized, hemisphere-specific process, that are susceptible to interference by a perceptual distractor. To further pinpoint neural mechanisms underlying this bias, we analyzed neural representations of the memory array and distractor with multivariate feature decoding based on the electrophysiological recordings.

### Fidelity of target and distractor neural representations govern mnemonic bias

To better understand the neural basis of the perceptual distractor bias, we computed the association between the neural representations of the target memoranda and the distractor, and the behavioral bias. Specifically, we hypothesized that the strength of encoding and maintenance of the memorandum would render it resilient to distractor bias whereas, conversely, the strength of encoding of the distractor would render it more potent at biasing responses.

For this, first, we performed trial-wise decoding of the memoranda orientations from time-resolved EEG activity using a multivariate decoder based on the pair-wise Mahalanobis distance^29,30^. The orientation of both memory array gratings could be decoded robustly from the (respective) contralateral electrodes shortly following their onset (Fig. 2A, 2B, left) (p<0.001, cluster-forming threshold p<0.05; n=23 participants for whom EEG data was available, see Methods). In addition, the orientations of both the cued and the uncued memoranda could be decoded for a few hundred milliseconds following the memory array onset (Fig. 2B, right; earliest distractor onset: 1.2 s), with the cued memorandum exhibiting significant decodability later into the maintenance epoch as compared to the uncued memorandum (Fig. 2B, inset, violin plots).

**Figure 2.**
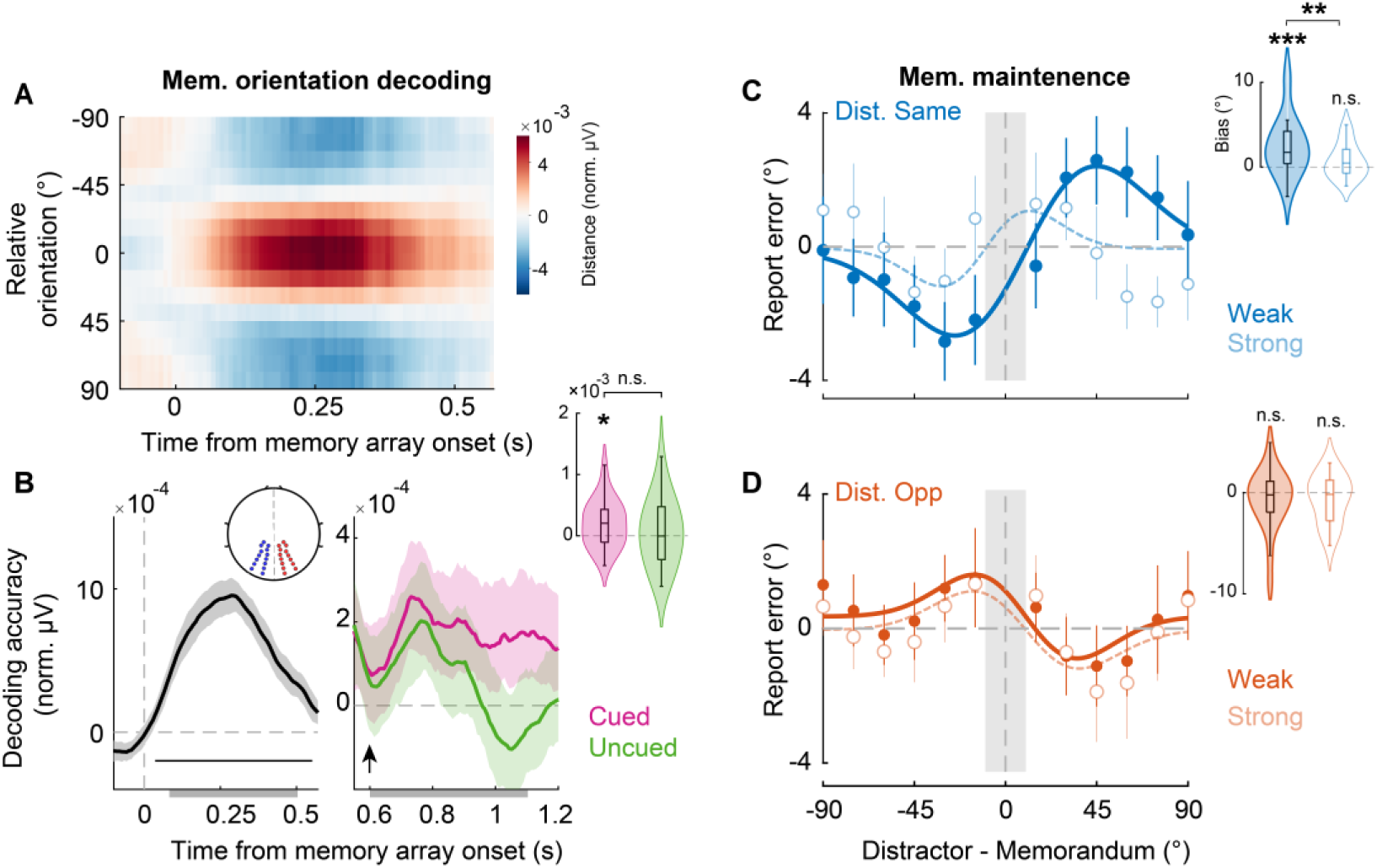
Stronger mnemonic neural representation mitigates distractor-induced bias. **A.** Orientation tuning map for memory array decoding for each time point relative to memory array onset. The tuning is computed as the similarity (sign reversed, mean-centered Mahalanobis distance) between the neural activity of the test trial and reference trials for stimuli at different relative orientations (y-axis). (All panels) Unless indicated otherwise, data averaged across participants (n=23, EEG) and for the left and the right gratings. **B.** (*Left*) Memoranda decoding accuracy (dot product of tuning curve with cosine, normalized uV, Methods) for each time point locked to the memory array onset (dashed vertical bar). Horizontal black bar: significant decoding epochs (cluster-based permutation test, Methods). Shading: s.e.m. Profiles were smoothed with an 80 ms rectangular moving window for clarity of visualization. *(Inset)* Posterior electrodes used for the decoding analysis, contralateral to the respective grating. (*Right*) Same as in the left panel but showing decoding accuracy separately for cued (magenta) and uncued (green) gratings following retro-cue onset (arrowhead). (*Inset*) Violin plots showing the distribution of average decoding accuracy across participants (n=23) for cued and uncued gratings following the cue onset until the earliest distractor onset (1.2s). **C.** Behavioral bias curves for the cued memorandum, but plotted based on a median split of memorandum maintenance strength – quantified with neural decoding accuracy during the early delay period (gray shaded bar, panel B, right) – when the distractor appeared on the same hemifield as the memorandum (n=23). Solid curve, filled symbols and filled violin plot (*inset*): Weaker maintenance trials. Dashed curve, open symbols and open violin plot (*inset*). Stronger maintenance trials. Other conventions are the same as in Figure 1F. **D.** Same as in C, but when the distractor appeared on the hemifield opposite to the cued memorandum. Other conventions are the same as in panel C and Figure 1G. (All panels) *: p<0.05; **: p<0.01; ***:p<0.001; n.s.: not significant.

Next, to test our hypotheses regarding the association between the fidelity of these neural representations and the distractor bias, we performed two broad types of median split analyses.

First, we performed a median split of trials into weak and strong “maintenance” – based on the strength of cued memorandum’s decoding accuracy (Methods) in a 500 ms window in the the delay period – beginning with cue onset, until 100 ms preceding the earliest distractor onset (Fig. 2B, right, gray shaded bar). We observed strong evidence for attractive bias – towards the distractor’s orientation – on the trials with weak memorandum maintenance, when the distractor appeared on the same hemifield as the cued memorandum (weak maint.: Bias_DistSame_=2.40±0.71, p<0.001, BF_+0_=29.5). By contrast, trials with strong memorandum maintenance exhibited no statistically significant evidence for an attractive distractor-induced bias (strong maint.: Bias_DistSame_=0.69±0.43, p=0.062, BF_+0_=1.26). Moreover, the attractive bias for the weak memorandum maintenance trials was significantly higher as compared to that for the strong maintenance trials (weak vs. strong maint.: p=0.009, BF_+0_=5.09, Fig. 2C). We also performed the same median split analysis for when the distractor appeared on the hemifield opposite to the cued memorandum, but did not find significant evidence for a repulsive bias modulation with maintenance strength (p>0.05 for all tests, Fig. 2D).

Additionally, we tested if the “encoding” strength of the cued memorandum also predicted a similar resilience to distractor bias. For this, we performed a median split analysis based on the decodability of the memorandum on the cued hemifield, but before the cue had appeared, in a window from 100 ms to 500 ms, following memory array onset (Fig. 2B, left, gray shaded bar). In contrast to maintenance strength, encoding strength did not modulate distractor-induced bias significantly. Specifically, we observed no statistically significant difference of cued memorandum encoding strength on bias either when the distractor appeared on the same hemifield as the cued memorandum (weak vs. strong enc.: p=0.78, BF_+0_=0.132), or when it appeared on the opposite hemifield (p=0.506, BF_-0_=0.39) (SI Fig. S3). Lastly, we tested if the distractor-induced bias for the cued memorandum would be influenced by the maintenance strength of the uncued memorandum; the latter was quantified, again, based on its neural decodability in a 500 ms window following cue onset (Fig. 2B, right). In this case also, we observed no significant effects of uncued memorandum maintenance strength on the bias either when the distractor appeared on the same hemifield (uncued-weak vs. strong maint.: p=0.763, BF_+0_=0.11), or on the opposite hemifield (p=0.052, BF_-0_=1.52), as the cued memorandum.

In other words, the robustness of maintenance, but not the strength of encoding, provided a clear predictor of susceptibility to distractor-induced bias, an effect that was strongly evidenced particularly when the distractor appeared on the same hemifield as the cued memorandum.

Second, we tested whether the strength of the perceptual distractor’s encoding influences the behavioral bias. For this, first, we decoded the orientation of the distractor from EEG traces locked to its onset. The distractor’s orientation could be decoded reliably for a few 100 ms, following its onset (p<0.001, cluster-forming threshold p<0.05) (Fig. 3A-B). Moreover, the strength of distractor encoding was not significantly different between strong and weak cued memorandum maintenance trials (p>0.4; Methods). As a result, any effects of distractor encoding strength on behavioral bias cannot be trivially explained by the aforementioned effects of cued memorandum maintenance.

**Figure 3.**
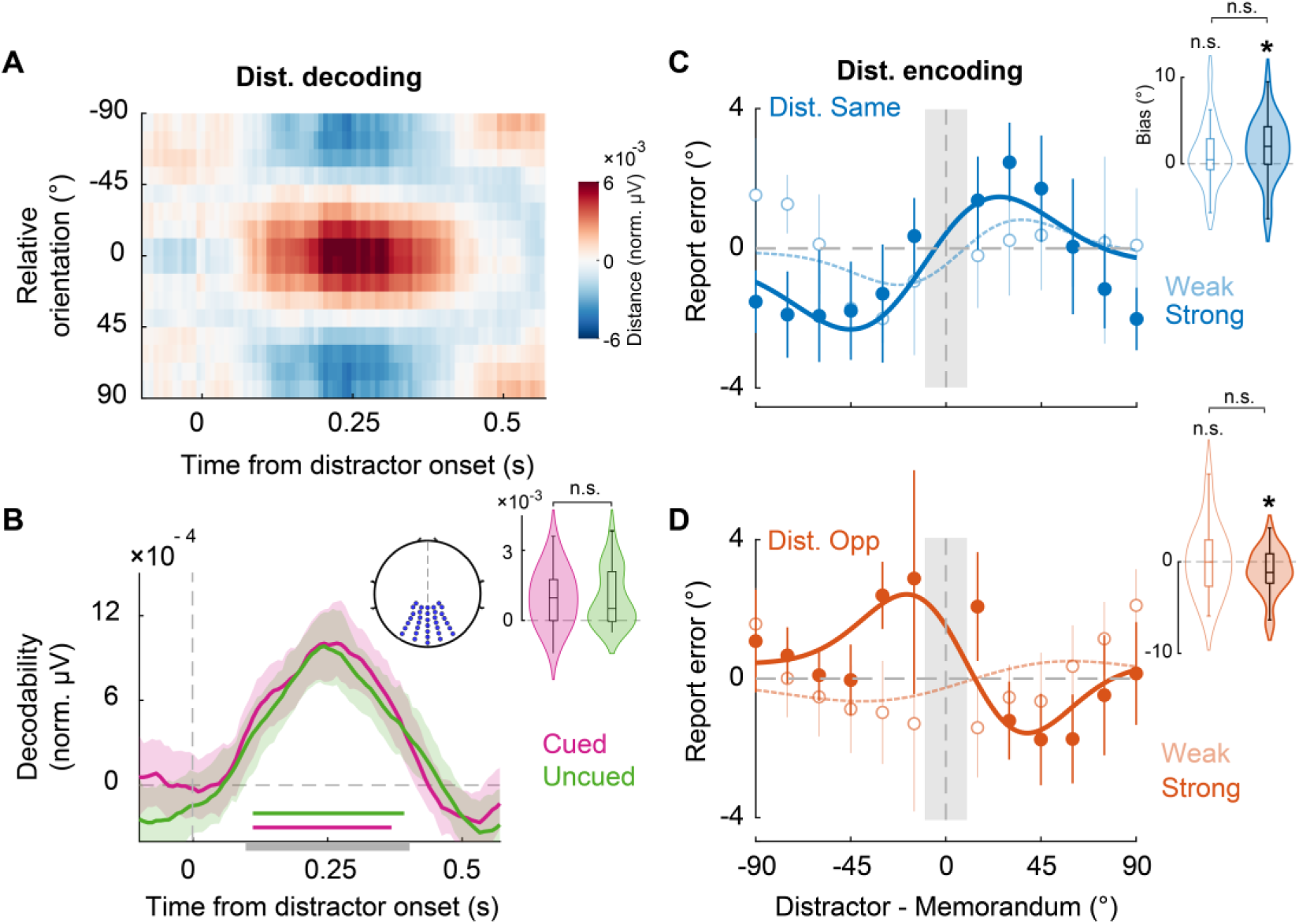
Stronger distractor encoding enhances distractor-induced bias. **A.** Same as in Figure 2A, but showing the orientation tuning map for distractor decoding for each time point relative to distractor onset. Data averaged across participants (n=23, EEG). Other conventions are the same as Figure 2A. **B.** Same as in Figure 2B (right), but showing distractor decoding accuracy for each time point locked to the distractor onset (dashed vertical bar). Magenta and green: Distractor decoding accuracy when it appeared on the cued or uncued hemifields, respectively. Horizontal bars: significant decoding epochs (cluster-based permutation test, Methods). Shading: s.e.m. *(Inset, top)* Posterior electrodes used for the decoding analysis. (*Inset, right*) Violin plots of distractor decoding accuracy on the cued (magenta) and uncued (green) hemifields. **C.** Same as in Figure 2C, but showing behavioral bias curves for the cued memorandum based on a median split of distractor encoding strength – quantified with neural decoding accuracy following distractor presentation (gray shaded bar, panel B) – when the distractor appeared in the same hemifield as the memorandum (n=23). Solid curve and filled violin plot (*inset*): Stronger distractor encoding trials. Dashed curve and open violin plot (*inset*). Weaker distractor encoding trials. Other conventions are the same as in Figure 2C. **D.** Same as in C, but when the distractor appeared in the hemifield opposite to the cued memorandum. Other conventions are the same as in panel C and Figure 2D. (All panels) *: p<0.05; **: p<0.01; ***:p<0.001; n.s.: not significant.

As before, for this analysis, we divided trials based on distractor’s orientation decoding accuracy in a window from 100-400 ms following its onset (Fig. 3B, gray shaded bar); we term these the “weak” and “strong” distractor encoding trials. When the distractor appeared in the same hemifield as the cued memorandum, distractor-induced bias was significantly attractive for strong distractor encoding trials (strong dist. enc.: behavioral Bias_DistSame_=1.79°±0.74°, p=0.006, BF_+0_=4.70. By contrast, this bias was not significantly different from zero for weak distractor encoding trials (weak dist. enc.: behavioral Bias_DistSame_=0.96°±0.75°, p=0.110, BF_+0_=0.80) (Fig. 3C). Similarly, when the distractor appeared in the hemifield opposite to the cued memorandum the distractor-induced bias was significantly repulsive on strong distractor encoding trials (Bias_DistOpp_=-1.1±0.53, p=0.022, BF_+0_=2.54, but not significantly different from zero for weak distractor encoding trials (Bias_DistOpp_=0.15±0.78, p=0.6, BF_-0_=0.189) (Fig. 3D). Even though these effects were individually significant, we did not observer a significant difference between the strong and weak distractor encoding trials for either relative location of the distractors (same or opposite hemifield as the cued memorandum) (Fig. 3C-D, insets); in this case, bias estimates were perhaps rendered noisier with only half of the number of trials within each split. Nevertheless, distractor encoding strength clearly modulated the magnitude of the antagonistic biases observed in behavior.

Taken together, these results reveal a mechanistic neural basis of perceptual distractor biases in visual WM. Stronger maintenance of the cued memorandum, as quantified by neural decodability during the delay period, produced weaker distractor-induced biases in behavioral reports. Interestingly, this bias was unaffected by memorandum encoding strength. Conversely, stronger encoding of the distractor produced stronger, antagonistic biases. We sought to identify additional task and neural variables that controlled the magnitude of the distractor bias.

### Distractor delay-period timing and gating control bias magnitude

To further understand the mechanistic basis of these distractor-induced biases, we investigated two specific factors that could affect the strength of these biases in WM. Based on recent literature^8^, we hypothesized that i) distractor’s timing of appearance – early versus late – during the delay period, and ii) its input gating – the extent to which the distractor is gated from access to WM – during the delay-period would modulate distractor-induced bias.

While distractor timing is readily quantified, to quantify distractor input gating we leveraged the finding that information on the prioritized (cued) hemifield is shielded more strongly against distractor interference (Fig. 2B, right)^56–58^. We explored neural signatures associated with the distractor that exhibited marked hemifield asymmetries between the cued and uncued hemifields. Interestingly, the strength of distractor encoding, as quantified by neural decodability over occipital cortex, was not significantly different regardless of whether the distractor appeared on the cued or the uncued hemifields (Fig. 3B) (decoding accuracy: Cued=9.9±2.6×10^−4^, Uncued=9.8±2.60×10^−4^, p=0.98, BF=0.22); thus, sensory decodability did not emerge as a reliable marker of distractor gating during the delay period.

We quantified distractor-evoked event-related potential (ERP) components, N1 and P2/P3a (Methods), which are known to index the allocation of spatial attention and working memory, respectively^31–33^. Although the distractor-evoked contralateral N1 amplitude was not different between hemifields (Fig. 4A) (Cued=-0.85±0.22, Uncued=-0.84±0.21, p=0.73, BF=0.22) the contralateral P2/P3a amplitude was significantly higher for the cued than the uncued hemifield (Cued=0.91±0.19, Uncued=0.72±0.18, p=0.015, BF=4.135). Consequently, we quantified P2/P3a amplitude as a potential indicator of distractor input gating (see Discussion for literature relevant to this claim). We then tested whether the distractor’s timing and input gating, as quantified by P2/P3 amplitude, would affect the distractor-induced bias in behavioral reports.

**Figure 4.**
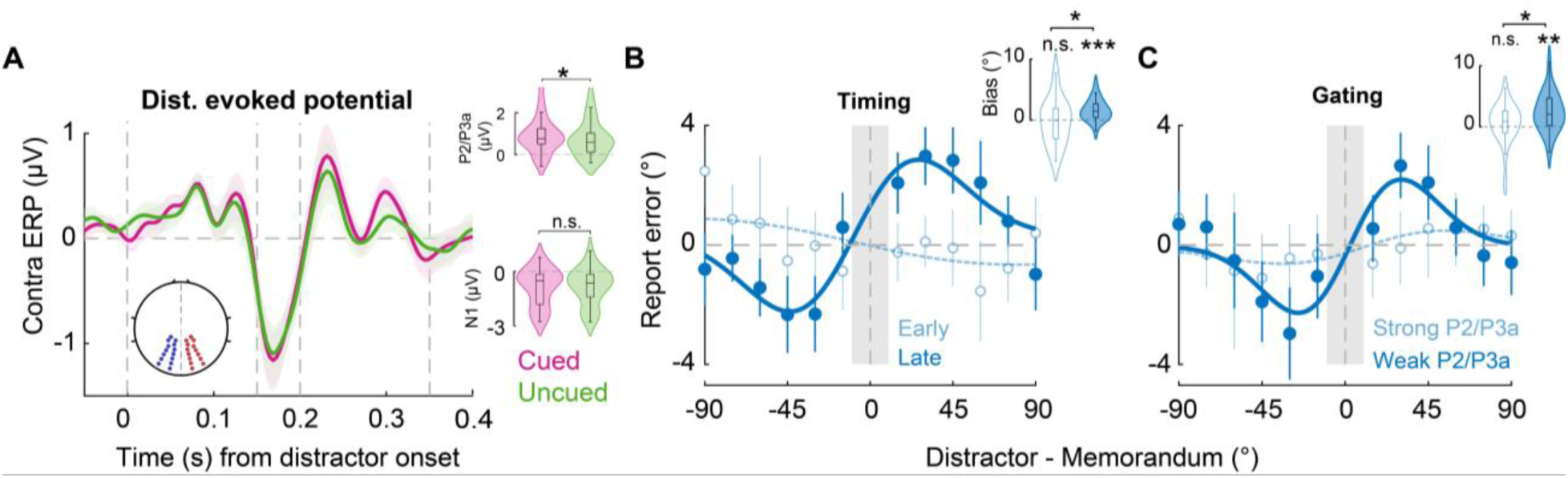
Distractor’s timing and gating predict mnemonic biases. **A.** Event-related potentials (ERP) locked to distractor onset (dashed vertical line at 0s) over posterior electrodes contralateral to distractor. Magenta and green traces: ERPs evoked by distractor in the cued and the uncued hemifields, respectively. Solid vertical lines at 0.15s-0.2s and 0.2s-0.35s demarcate the temporal windows for computing N1 and P2/P3a amplitudes, respectively. Shading: s.e.m. (*Insets*) Distribution of the P2/P3a (*top*) and N1 (*bottom*) amplitudes evoked by the distractor in the cued (magenta) and uncued (green) hemifields, respectively. **B.** Same as in Figure 3C, but showing behavioral bias curves for the cued memorandum separately for trials in which the distractor appeared early (dashed, open symbols) or late (solid, filled symbols) during the delay period, when the distractor appeared on the same hemifield as the memorandum. (*Inset*) Violin plots show the distribution of bias strengths across participants (n=23) for early (open) and late (filled) distractor trials. **C.** Same as in panel B, but showing behavioral bias curves for the cued memorandum separately for trials with strong (dashed, open symbols) or weak (solid, filled symbols) distractor-evoked P2/P3a amplitude, a putative marker of distractor gating (panel A). (*Inset*) Violin plots showing the distribution of bias strengths across participants (n=23) for strong (open) and weak (filled) distractor gating trials. (All panels) *: p<0.05; **: p<0.01; ***: p<0.001; n.s.: not significant.

For the first analysis, we median split trials based on when the perceptual distractor appeared during the working memory epoch; we term these “early” and “late” distractor trials; as with all previous analyses, these also included only the trials in which the cued grating was probed. Because our task design incorporated a fixed delay between the memory array and response probe presentation, early (late) distractors appeared earlier (later) during the WM maintenance epoch, and were temporally more distal (proximal) to the response probe (Fig. 1A). Distractor-induced bias, when the distractor appeared on the same hemifield as the cued grating, was significantly positive for late distractor trials (“late”: behavioral Bias_DistSame_=1.6°±0.4°, p<0.001, BF_+0_>10^2^) (Fig. 4B). By contrast, this bias was not significantly different from zero for early distractor trials (“early”: behavioral Bias_DistSame_=-0.3°±0.8°, p=0.64, BF_+0_=0.16) (Fig. 4B). Moreover, the bias was significantly stronger for late distractors compared to early distractor trials (p=0.014, BF=3.4).

For the second analysis, we divided trials based on the amplitude of the perceptual distractor-evoked contralateral P2/P3a component, for trials in which distractor appeared on the same hemifield as the cued grating. Distractor-induced bias was not significantly different from zero for high P2/P3a-amplitude trials (“strong”: behavioral Bias_DistSame_=0.63°±0.68°, p=0.20, BF_+0_=0.52) (Fig. 4C). By contrast, this bias was significant, and attractive, for weak P2/P3a amplitude trials (“weak”: behavioral Bias_DistSame_=2.4°±0.66°, p<0.001, BF_+0_=47.03) (Fig. 4C). Moreover, we observed a significant difference between the high and low P2/P3a amplitude trials, such that putatively weakly gated trials revealed a stronger bias than strongly gated trials (p=0.030, BF=1.90).

We repeated the same median split analyses for trials in which the distractor appeared on the hemifield opposite to the probed memorandum. In this case, while qualitative trends were apparent in the repulsive bias along expected lines – stronger biases for late and weakly gated distractor trials – none of these trends were statistically significant (SI Fig. S4A-B).

Finally, we report an omnibus analysis combining across all three perceptual distractor dependent factors: encoding strength, timing and gating. For this, we first tested and confirmed that these three factors were not mutually correlated with each other. We found no significant difference in the strength of distractor encoding between early and late distractor trials (p=0.153, BF=1.13). Similarly, there was no significant difference in distractor-evoked P2/P3a amplitude between early and late distractor trials (p=0.563, BF=0.22). Finally, there was also no significant difference in P2/P3a amplitude between trials with strong versus weak distractor encoding (p=0.362, BF=0.37). Given their trail-wise independence, we pooled together trials across functionally matched levels of all three factors (SI Fig. S4C-D) – strong encoding strength, late timing and weak gating (“strong interference”) – as well as their converse counterparts – weak encoding strength, early timing and strong gating (“weak interference”). In the strong interference trials, we observed robust evidence for antagonistic biases: a strongly attractive bias (Bias_DistSame_=1.94°±0.35°, p<0.001, BF_+0_> 10^4^) or a repulsive bias (Bias_DistOpp=_=-0.77°±0.37°, p=0.02, BF_-0_=1.96), when the distractor was on the same or opposite hemifield as the cued memorandum, respectively. By contrast, no such trends were apparent in the weak interference trials (Bias_DistSame_=0.4°±0.43°, p=0.181, BF_+0_=0.33; Bias_DistOpp_=-0.011°±0.34°, p=0.486, BF_-0_=0.14).

Overall, these results show that perceptual distractors that appeared later during working memory maintenance, produced a stronger bias in orientation reports of the cued memorandum. Moreover, the effectiveness of distractor input gating – as quantified operationally by the distractor-evoked P2/P3a amplitude – reliably predicted the magnitude of bias. A combination of these factors – stronger distractor encoding, late timing and weak gating – produced robust, antagonistic patterns of distractor-induced bias in behavioral responses. We synthesized these links between neural and behavioral effects with a computational attractor model of visual WM.

### An attractor model explains distractor-induced biases in visual WM

To provide a parsimonious, mechanistic explanation of these perceptual distractor-induced biases during the delay period, we designed a two-tier bump attractor model. Based on extensive recent literature^3–5,9,13,18,34–38^, working memory maintenance in our model involves both the sensory (visual) cortex (VC), and higher cortex (HC), such as the prefrontal cortex (PFC) or the parietal cortex (PPC). In fact, the latter regions both are reported to be involved in robust maintenance in the presence of distractors^9,23,24,39^. Moreover, a key element of novelty in our model is as follows: while distinct attractors independently encode stimuli in the two hemifields, they interact with each other via competitive, cross-hemifield inhibition, based on biological observations^40–44^. We provide a brief description of the model below; a detailed description is provided in the Methods.

The VC and the HC were modeled with separate ring attractors; we also employed distinct attractors for encoding the memorandum in each (cued, uncued) hemifield, yielding a total of 4 attractor networks (Fig. 5A). Each attractor network comprised both excitatory (E) and inhibitory (I) neurons (512 each) with dynamics described by rate-based models (Methods, equations 1-2). Recurrent connections among E neurons within an attractor were local and topographic (Fig. 5A, top), whereas all other connection types (E-I, I-E, I-I) were global (see Methods). VC neurons provided input to HC neurons via topographic, excitatory feedforward connections and received input from HC via reciprocal, topographic feedback connections (Methods, equations 3-4). Based on our experimental observation, we expected cued information in the VC to be more robust to distractors. Therefore, HC→VC feedback connections were 20% stronger for the cued, than the uncued, hemifield (Fig. 5A, thick HC-VC connection). In addition, previous studies suggest that mnemonic information in higher order areas (e.g., PFC, PPC) is relatively robust to distractors^8,9,13,23,39^. Consequently, we transiently blocked feedforward connections from the VC to the HC during the maintenance epoch, thereby rendering the HC mnemonic information immune to disruption by the distractor upon its presentation; this input was selectively reinstated for the weak distractor input gating experiments, described below. Finally, based on previous reports of competitive visual inhibition across the hemifield^40–44^, we also modeled cross-hemispheric inhibitory connections between the VC attractors in each hemisphere (Fig. 5A, black circles; equation 3); these topographic connections played a critical role in explaining the antagonistic pattern of biases (see next).

**Figure 5.**
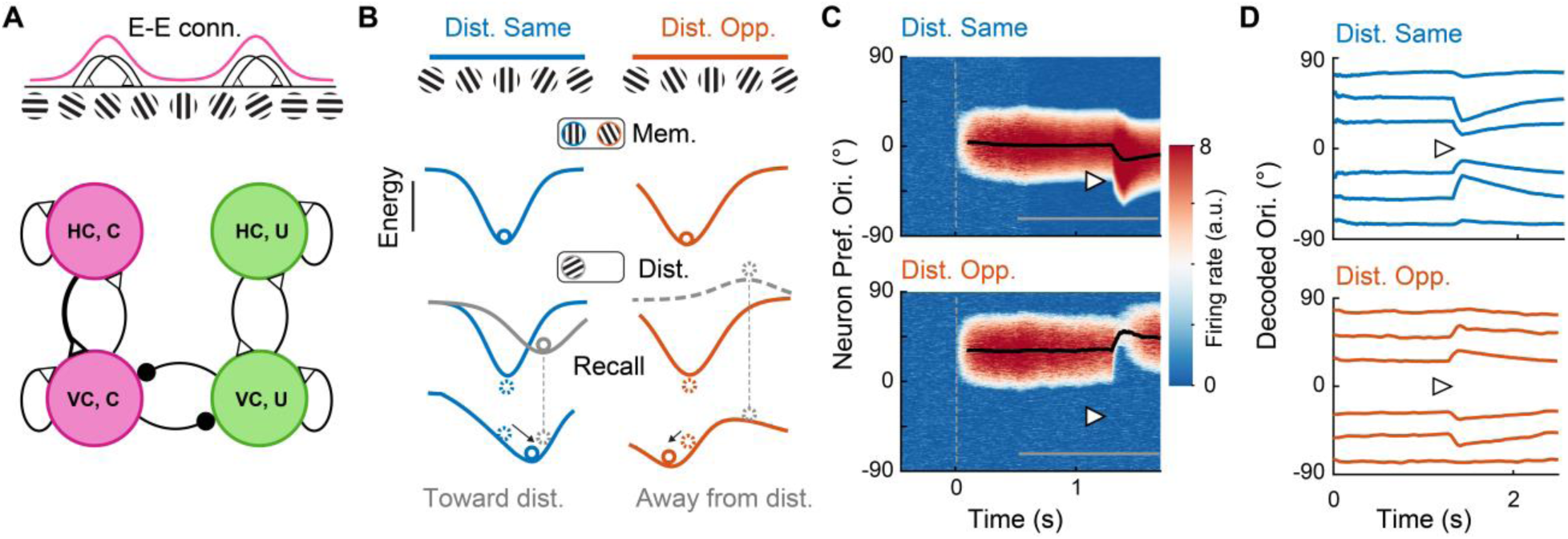
Two-tier attractor model explains spatially antagonistic, mnemonic biases. **A.** Two-tier ring attractor model. The visual cortex (VC, bottom) and higher cortex (HC, top) were modeled with separate ring attractors. Distinct attractors also encoded stimuli in the cued (C, left, magenta) and the uncued (U, right, green) hemifields. Top-down input from HC to VC was stronger for the cued (thick black line) compared to the uncued hemifield. Mutual inhibition between the VC,C and VC,U attractors was mediated by topographic, cross-hemispheric inhibitory connections (black circles with horizontal arcs). (*Inset, top*) Recurrent excitatory (E-E) connectivity schematic showing local and topographic connectivity profile (see text for details). **B.** An energy landscape intuition for spatially antagonistic distractor-induced biases. (*Top row*) The memorandum’s orientation (blue or orange circle) is maintained at the lowest energy state (highest firing rate). (*Middle row, left*) When the distractor (gray circle) appears in the same hemifield as the memorandum, it causes a dip (energy minimum; solid gray line) in the energy landscape. (*Middle row, right*) By contrast when the distractor appears in the opposite hemifield, it causes a peak (energy maximum; dashed gray line) in the energy landscape. (*Bottom row*) As a result, the final energy landscape, obtained by summing the memorandum and distractor energies, attracts the memorandum’s representation towards the distractor’s orientation when the latter appears in the same hemifield (blue circle), whereas it repels the memorandum’s representation away from the distractor’s orientation when the latter appears in the opposite hemifield (orange circle). **C.** Firing rate heat maps of the neural activity; hotter colors indicate higher activity. y-axis: E neurons arranged in order of orientation preference on the ring; x-axis: Time. (*Top)* In this example, the onset (dashed vertical line) of the memorandum gratings (0° orientation) causes a localized bump in firing rate at the respective location on the ring, which persists throughout the delay period (gray horizontal bar). Thick black curve: orientation decoded from the population activity vector. When the distractor grating (black triangle; −30° orientation) is presented transiently, and in the same hemifield as the memorandum, the latter’s representation is biased toward the distractor’s orientation. *(Bottom)* Same as in the top panel, but in this case the distractor grating (−30° orientation) is presented transiently, and in the opposite hemifield as the memorandum (+30° orientation). Here, the memorandum’s representation is biased away from the distractor’s orientation. **D.** Decoded neural representations across time for six different orientations of the memoranda (curves, distinct simulations). Distractor grating (black triangle; 0° orientation) presented in the same (*top,* blue curves) or the opposite (*bottom,* red curves) hemifield as the respective memoranda.

With this model, we sought to replicate the attractive and repulsive biases that occurred across space. We explain the expected sequence of events, first, with an energy landscape model (Fig. 5B); detailed simulations are presented subsequently. The energy landscape provides schematic representation activity and attractors in the network^45–47^: Minima (valleys) in the energy landscape reflect stable fixed points that cause stored patterns to drift towards their location whereas, conversely, maxima (peaks) reflect unstable fixed points that cause stored patterns to drift away from them.

The presentation of a memorandum produces a valley in the VC attractor’s energy landscape at its orientation, which persists during the maintenance epoch due to recurrent excitation (Fig. 5B, top). Next, the distractor appearance in the same hemifield as the memorandum (DS) causes another dip in the VC attractor’s energy landscape at the distractor’s orientation (Fig. 5B, left column, middle row, gray trace). If the dip induced by the distractor is large enough, it perturbs the minimum in the landscape causing the minimum to shift, and the maintained representation to drift, towards the distractor’s orientation (Fig. 5B, left column, bottom row, blue trace); this yields an attractive bias in the distractor’s hemifield (Fig. 5B, left column, bottom row, blue circle). By contrast, the distractor also creates a peak at its own orientation in the VC attractor’s energy landscape in the opposite hemifield (DO); this energy peak arises as a result of the topographic, cross-hemispheric inhibitory connections (Fig. 5B, right column, middle row, gray trace). As schematized, this peak repels the minimum in the landscape, and causes the maintained representation of the memorandum to drift away from the distractor’s orientation (Fig. 5B, right column, bottom row, orange trace); this yields a repulsive bias in the hemifield opposite the distractor (Fig. 5B, right column, bottom row, orange circle).

Next, we describe the results of simulating the model (parameters in SI Table S1). We modeled the initial memory array presentation with an additive input current to the VC-E neurons of each attractor (cued, uncued) using a von Mises spatial profile centered on the orientation of the respective grating (***κ***_*M*_ =2.7°, equation 5). This initiated localized bumps of activity in VC attractors, at locations corresponding to the orientations of the encoded gratings (Fig. 4C top and bottom, activity at 0° and +30°, respectively). Upon disappearance of the memory array, during the delay period (Fig. 5C, gray underbar), the bump of activity was maintained through recurrent excitation in each attractor^26,27,48^ (Fig. 5C, red patches). A distractor was presented at t=1300 ms during the delay period (Fig. 5C, triangle); distractor presentation was modeled, also with a von Mises input profile (***κ***_*D*_ =2.3°, equation 6) centered on the distractor’s orientation (Fig. 5C; distractor at −30° for both panels).

Upon its presentation the distractor induced antagonistic mnemonic bias across hemifields (Fig. 5C). When the distractor was presented in the same hemifield as the memorandum the bump of activity shifted systematically toward the distractor’s orientation (Fig. 5C, top, heat map). By contrast, when it was presented in the opposite hemifield as the memorandum the bump of activity shifted systematically away from the distractor’s orientation (Fig. 5C, bottom, heat map). We repeated these simulations with six different memoranda of different orientations, with a distractor always appearing at 0° orientation. In each case, the distractor biased the decoded neural representations towards (Fig. 5D, top) or away from (Fig. 5D, bottom) itself, albeit to varying degrees (see next).

Next, we sought to replicate key experimental observations regarding the effects of perceptual distractors on behavioral bias magnitude (Methods). Behavioral estimates of orientation were obtained by averaging the memorandum’s orientation decoded from both VC and HC neural activity at the end of the trial; this averaging rendered behavioral estimates more robust to noise (see SI Results Section for a detailed analysis of error-correcting dynamics).

First, we studied the effect of the distractor’s location on simulated behavioral biases. The distractor produced an attractive bias when it appeared in the same hemifield (Fig. 6A) and a repulsive bias when it appeared the opposite hemifield as the memorandum. As in the experiments, the magnitude of this bias varied parametrically with the angular difference between the distractor and the memoranda: the strongest bias appearing for intermediate angle differences, and the weakest appeared for extreme differences (0° or ±90°) (Fig. 1G-H). Neural bias – with orientation decoded based on the peak of the activity bump in the VC attractor (Methods) – revealed an identical trend (SI Fig. S5A).

**Figure 6.**
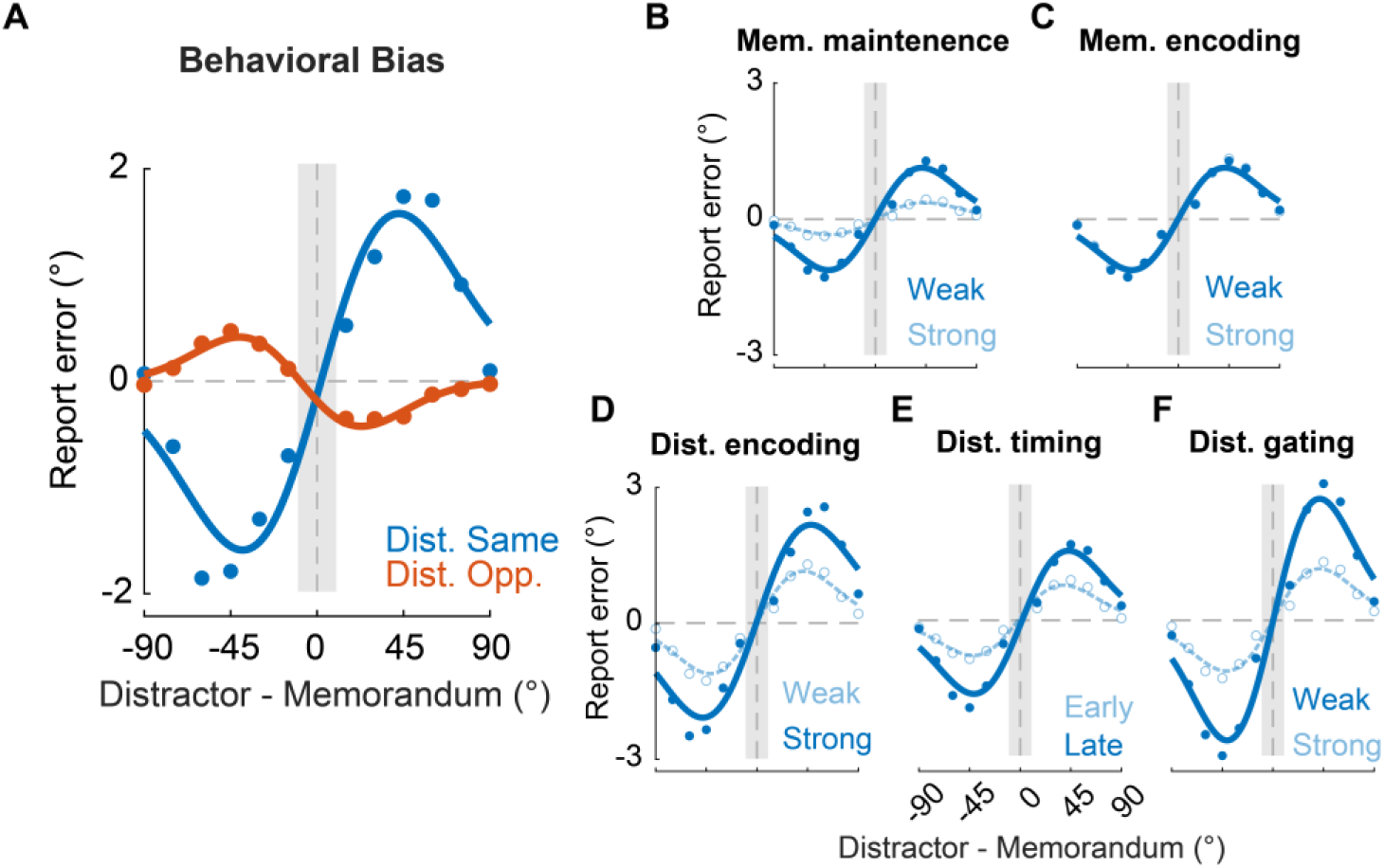
Model simulations qualitatively replicate distractor-induced mnemonic biases. **A.** The model’s simulated behavioral bias when the distractor appeared in the same (blue), or opposite (orange), hemifield as the cued memorandum. Compare with Figures 1F-G (experimental data). Other conventions are the same as in Figures 1F-G. **B.** Same as in panel A, but showing behavioral bias for the cued memorandum plotted separately for simulated trials with weak (solid curve and filled symbols) or strong (dashed curve and open symbols) memorandum maintenance trials (compare with experimental Figure 2C). **C.** Same as in panel B, but for simulated trials with weak (solid curve and filled symbols) or strong (dashed curve and open symbols) memorandum encoding. **D.** Same as in panel B, but for simulated trials with weak (dashed curve and open symbols) or strong (solid curve and filled symbols) distractor encoding (compare with experimental Figure 3C). **E.** Same as in panel B, but for simulated trials with early (dashed curve and open symbols) or late (solid curve and filled symbols) distractor presentation in the delay-period (compare with experimental Figure 4B). **F.** Same as in panel B, but for simulated trials with weak (solid curve and filled symbols) or strong (dashed curve and open symbols) distractor gating (compare with experimental Figure 4C) (B-F) Other conventions are the same as in Figure 2B. See text for details of how each simulation condition was controlled.

Second, we studied the effect of memorandum maintenance strength on distractor-induced biases (Fig. 6B). We modeled the strength of memorandum maintenance by modulating feedback input from the HC to VC by 2x; we hypothesized that stronger feedback would render mnemonic representations in the VC more robust to interference via error-correcting dynamics^8,9,24^. Indeed, stronger top-down feedback from the HC to VC enabled more accurate maintenance of the memorandum features during the delay period (see SI Results section on *Error-correcting dynamics in a two-tier attractor model* and SI Fig. S6). As a result, the magnitude of distractor-induced bias was systematically higher for the weakly maintained memoranda (Fig. 6B, solid), compared to the strongly maintained ones (Fig. 6B, dashed), consistent with experimental results. Interestingly, increasing the memorandum’s encoding strength – by affording a 60% higher input current amplitude to the memorandum during the encoding phase – by itself, did not modulate distractor bias (Fig. 6C), precisely in line with experimental observations.

Third, we studied the effect of distractor encoding on bias (Fig. 6D). Here, we modulated distractor encoding strength by affording a 60% higher input current amplitude to strongly, as compared to weakly, encoded distractors. The stronger perturbation of the energy landscape induced by the strongly encoded distractors yielded faster drifts in the memorandum’s representation thereby yielding a higher bias for strongly (Fig. 6E, solid) compared to weakly (Fig. 6E, dashed) encoded distractors; again, these results matched with the experimental observations.

Fourth, we found that late distractors produced stronger mnemonic biases (Fig. 6F). We propose that this occurred because of the timing of the distractor vis-a-vis the error-correcting dynamics via top-down feedback from HC to the VC. Specifically, late distractors appear later during the WM maintenance epoch and are temporally more proximal to the response probe than early distractors. Consequently, the former received ameliorative top-down feedback for a shorter duration than the latter. This produced a higher (residual) bias for late (Fig.6F, solid) compared to early (Fig. 6F, dashed) distractor trials at the end of the delay epoch, again, matching experimental findings.

Fifth, we studied the effect of distractor input gating on simulated biases (Fig. 6F). We modulated the extent of distractor gating by introducing feedforward input from the VC to HC (strength: 0.01) uniformly during the delay period. We hypothesized that stronger feedforward input into the HC (“weak input gating”) would render mnemonic representations in the HC more prone to interference. Indeed, the magnitude of distractor-induced bias was systematically higher for the weakly-gated distractors (Fig. 6F, solid), compared to the strongly gated distractors (Fig. 6F, dashed), consistent with experimental results (see also SI Fig. S5).

Finally, distractor effects were stronger (magnitude higher) when the distractor appeared in the same hemifield as the memorandum, as compared to when it appeared in the opposite hemifield, uniformly across all simulations (Fig. 6A, SI Fig. S5B-F). Notably, this disparity in bias magnitudes occurred also in the experimental data across hemifields (Fig. 1G-H). In our model this can be explained by the fact that the distractor’s influence in the hemifield opposite the memorandum, mediated by the indirect, cross-hemifield inhibition, decayed comparatively faster than its effect in the same hemifield, which was mediated by direct, local excitation.

Taken together, our results uncover space-specific, antagonistic patterns of induced by perceptual distractors in visual working memory. Behavioral reports were biased towards (away) the distractor’s orientation when it appeared in the same (opposite) hemifield as the memorandum. Cued memoranda were more strongly shielded from distractor interference as compared to uncued memoranda. A two-tier ring attractor model provided a mechanistic explanation for the origins of these biases. Moreover, the model comprehensively explained diverse factors that mediate distractor-induced bias, including cued memorandum maintenance, distractor encoding, timing in the delay period, and input gating. Overall, the results identify lateralized sensory buffers as the putative neural underpinnings of perceptual distractor-induced biases in visual working memory.

## Discussion

We discovered that a salient perceptual distractor that shares task-relevant features with the memoranda produces space-specific, antagonistic biases in visual working memory. Memorandum reports were biased towards the distractor’s orientation when the latter occurred in the same hemifield as the former, but were biased away when these stimuli appeared in opposite hemifields^22,49–52^. Over time, mnemonic information became more labile and prone to interference by distractors that appeared later in the maintenance epoch. Yet, information associated with the selected (retro-cued) hemifield was held more robustly and shielded against distractor interference, possibly by “error-correcting” top-down control mechanisms^8,9,13,19,24^. Nevertheless, we show that lateralized sensory buffers could enable task-irrelevant perceptual input – sharing features with the memoranda – to gain access to mnemonic information, and thereby, influence the contents of WM profoundly.

Several lines of evidence show that the perceptual distractor biased the information maintained in WM, rather than merely inducing a bias in the behavioral report (response bias). First, had the distractor merely produced a response bias, we would have expected a similar bias to occur regardless of the distractor’s location and its orientatation. In our task, even though the response probe and bar were always presented centrally, we observed distinct patterns of biases depending on where the distractor occurred relative to the probed memorandum, and in a manner that systematically varied with their relative orientations (Fig. 1G-H). Second, the distractor was flashed briefly (100 ms) and was followed immediately by a noise mask (pedestal) to diminish its persistent representation. Moreover, participants were aware that the distractor was task irrelevant, and it appeared at least 500 ms before the probe, rendering biases arising from persistence in iconic memory unlikely. Third, we performed a control analysis by fitting the participants’ responses based on a combination of target and distractor orientations (von Mises mixture model), to model the scenario where participants reported the distractor’s orientation on a subset of trials in which, perhaps, their memory of the memorandum orientation was weak. Formal model selection analysis, based on AICc and BIC, unequivocally ruled out this scenario.

Our results strongly support the sensory recruitment hypothesis of visual WM, which posits that brain regions involved in processing the memorandum during encoding also contribute to its maintenance in WM^1–5^. We were able to successfully decode the orientation of the memoranda during the delay epoch from occipital electrodes, and also showed that contralateral decoding strength was a key determinant of distractor-induced biases in behavioral reports (Fig. 2A-B). Yet, due to the low spatial resolution of EEG we cannot readily disambiguate whether the neural signatures that we observed originated entirely from the early visual cortex or also included contributions from other areas, like the parietal or prefrontal cortex^49^. Lastly, distractor-induced bias effects were hemifield-specific; this is consistent with previous studies which suggest that storage of mnemonic information involves a lateralized buffer including, possibly, also the sensory cortex ^22,50–53^. Such contralateral buffers may offer a conduit to perceptual distractors to bias the contents of working memory, especially when such information is labile to interference, such as during the delay-period.

Nevertheless, prioritization in working memory promoted distractor resilience: cued memoranda were better shielded from distractor interference than uncued memoranda. Taken together with previous studies^24,54^, our results show that prioritized information is more robust to interference, possibly due to feedback-mediated stabilization of the cued memorandum’s representation^24,54^. Indeed, when simulated in our computational model, top-down feedback from higher cortex stabilized WM representation in the visual cortex on the cued hemifield (Fig. 6B). On the other hand, it is also possible that such stabilization of the cued representation arises from a mixture of persistent activity and activity-silent states^7,29,48,55^; the latter are arguably less prone to interference from sensory distractors^7^. Future experiments can disambiguate these hypotheses by using a “pinging” protocol that activates latent representations and studying the effect of distractor interference on these activated representations.

Among the multiple electrophysiological indices we tested, we propose the P2/P3a ERP amplitude as a putative marker for distractor gating into working memory. Previous literature has often linked the P2 component to the gating (or suppression) of distractor information during working memory tasks^56–58^. For instance, higher distractor-evoked P2 amplitude, indicative of greater distractor gating, correlated positively with performance in an auditory WM task^31^. Similarly, the short peak latency P3 component, referred to as the P3a, possibly originates in the frontal lobe^57^ and indexes top-down inhibition of distractors^56,57.^ In fact, distractor-evoked P3a amplitudes were correlated with frontal lobe grey matter volume across individuals^59^, suggesting its involvement in the distractor-related processing. Our study extends these findings and shows that the P2/P3a amplitude contralateral to the distractor was higher at the cued compared to the uncued hemifield; in other words, the P2/P3a may index a stronger input gating of cued mnemonic information stored in a lateralized mnemonic buffer (Fig. 4A). Moreover, the P2/P3a amplitude directly predicted the magnitude of the distractor-induced bias, providing further confirmation for this “gating hypothesis”.

We explained the mechanistic origins of perceptual distractor biases with a two-tier ring attractor WM model. Attractor models have been frequently used for explaining diverse phenomena in WM, including variable precision^27^, serial biases^48^, categorical biases^60^ and the like^28^. For instance, a recent study employed an attractor model to explain serial biases in spatial WM^48^. Other studies employing attractor models have been able to explain the mechanisms behind trial to trial variability and the presence of categorical biases in visual WM^26,27,48^. In our study, a ring attractor model was a natural candidate to explain our results. Recent research has shown that maintenance in visual working memory (WM) involves multiple regions across the brain, including both sensory and higher-order cortex^18,35,37^. We, therefore, employed a two-tier attractor model that includes both sensory and higher order brain regions to simulate our experimental findings.

In our model, a higher-order cortical region (HC) stored information relevant to the memorandum and provided top-down influence to the visual cortex (VC) during the delay period to mitigate distractor interference in WM. Candidate regions for the HC include parietal cortex or prefrontal cortex both of which are known to contain robust WM representations, especially in the presence of distractors^9,13,23,24,39^. In our model, the representation in HC was rendered immune to distractor interference by transiently blocking its bottom-up input from VC during the delay period, and enabling this input only for weakly gated distractors. Neurophysiological mechanisms that could mediate such dynamic connectivity include oscillatory coherence^61^ or neuromodulatory gating (e.g., acetylcholine^62^). Yet, some recent studies have shown that disruption in visual WM maintenance occurs in both lower (V1, V4) and higher level cortex^11^. In this case, the coordinated activity across these regions could help improve the robustness of maintenance in WM, even in the presence of distractors^18,35–37,63^. Consistent with these results, our simulations showed that combining VC and HC activity yielded a more accurate behavioral readout of the memorandum’s orientation, as compared to either VC or HC activity alone (SI Fig. S5A). Future work, employing fMRI, perhaps in combination with EEG, could help identify the precise brain regions, and their dynamics, that mediate distractor biases in WM.

Our model accurately reproduced the effect of distractor timing on mnemonic bias: distractors presented later during the delay epoch induced a stronger bias, and vice versa. This experimental result is in line with literature suggesting that neural WM representations are not static but evolve over time, as dynamic processes^8,30,64^. We leveraged these findings in our model to explain distractor timing effects. In our model, the bias due to error-correcting dynamics in visual WM decays steadily; these dynamics are, in turn, mediated by top-down feedback from higher cortex (e.g., PFC^9,13,23,24,39^). Due to this effect, early distractor trials provided more time for the error-correcting mechanism to operate thereby yielding a weaker behavioral bias, as compared to late distractor trials. While model simulations accurately recapitulated experimental data, future experiments can test this hypothesis by stimulating neurons in higher brain regions during WM maintenance and studying its effects on the extent of error-correcting dynamics observed in lower order brain regions and in behavior.

In conclusion, our study offers putative mechanistic insights into neural processes by which perceptual distractors induce space-specific biases in working memory. Complementing these experimental results, our simulations suggest key computational hypotheses that by which distractors bias in WM representations. These hypotheses may be tested causally with non-invasive brain stimulation (e.g., transcranial magnetic stimulation) by focally perturbing activity in specific brain regions (e.g., visual cortex, prefrontal cortex), and at specific times during the delay epoch. Understanding the neural basis of distractor interference could have critical implications for improving the robustness of working memory in patients with attention deficit disorder (ADD) or mild cognitive impairment (MCI^65^).

## Materials and Methods

### Participants

24 participants (7 female; age range: 23-38 yrs; median age: 28 yrs), all right-handed, participated in the experiments. All participants had normal, or corrected-to-normal, vision and no known history of neurological disorders. Participants provided written and informed consent to take part in the experiments. Study protocols were approved by the Institute Human Ethics Committee at the Indian Institute of Science, Bangalore.

#### Sample size estimation

Sample sizes were not estimated with power analysis. The sample size in this study (n=24) is largely similar to that of several earlier studies on working memory and EEG-based grating orientation decoding^12,29,30^.

### Behavioral data acquisition

#### Task and stimuli

Participants completed the experimental tasks in a dimly lit room while seated, with their heads secured by a chin rest, with the eyes located 60 cm from a stimulus display screen. Stimuli were displayed on a contrast calibrated (i1 Spectrophotometer, X-Rite Inc.) 24-inch LED monitor (XL2411Z, BenQ corp.) with a resolution of 1920×1080 pixels and a refresh rate of 100 Hz. Stimuli were programmed in Matlab 2015b (Mathworks Inc.) with Psychtoolbox version 3.0.15. During the experiment, participants used an optical mouse (M-U0026, Logitech) to provide responses. Participants’ gaze position was tracked throughout the experiment using an infrared eyetracker (500 Hz, HiSpeed, iView X, SensoMotoric Instruments Inc.).

Each trial began with a fixation dot (full black, radius: 0.3 dva) presented at the center of the screen. Concurrently with this, circular placeholders (uniform white noise patches) were presented symmetrically in either hemifield along the azimuth (eccentricity: ±8.0 dva, radius: 4.0 dva); both the fixation dot and the placeholders remained on the screen throughout the trial. 500 ms after fixation dot onset, the memory array, comprising two sine-wave gratings (50% contrast, radius: 3.5 dva, spatial frequency: 0.9 cpd), appeared for a brief interval (100 ms), each embedded (50% alpha blended) concentrically inside one of the placeholders in each hemifield. 500 ms after memory array offset a retro-cue (white filled semicircle) appeared for 100 ms which indicated the location of the grating that would likely be probed for response (cue validity: 70%). Following a random delay (500-1250 ms, drawn from an exponential distribution), a singleton distractor grating (100% contrast) appeared for 100 ms in one of the two hemifields, pseudorandomly, with equal probability. This “perceptual” distractor’s size and spatial frequency was identical to that of the memory array gratings, but its orientation was drawn randomly from a uniform distribution across all angles at least ±10° away from either grating’s orientation in the memory array. The distractor was irrelevant to the task and the participants were explicitly instructed about the same. No distractor grating appeared in 20% of the trials; these trials interleaved among the distractor presentation trials.

A fixed interval (1950 ms) after retro cue onset, a response probe (white or black filled circle, 0.3 dva, with a yellow ring around it) appeared on the screen indicating the grating relevant for response. A white (black) probe indicated that the participants must reproduce the orientation of the cued (uncued) grating. A randomly oriented bar (length: 7 dva, width 0.3 dva) appeared at the center of the screen upon response initiation. Participants reproduced the orientation of the grating on the probed hemifield from memory by rotating the bar with the mouse using their right (dominant) hand, and pressed the left mouse button to register their response. Although there was no constraint on the time to initiate the response after probe onset, participants had to complete their response, once initiated, within 2 seconds. The next trial began after an inter-trial interval of 2 seconds.

#### Training and testing

Before the main experiment, participants completed between 2-4 blocks (100 trials each) of the task, typically, without EEG recordings. To familiarize and train participants on the task, feedback was provided at the end of each trial. The feedback consisted of an oriented, magenta bar whose orientation precisely matched that of the probed grating shown along with the reported orientation (black bar); the feedback was shown for 200 ms at the end of the response period. Following this, they also received textual feedback, displayed on the screen, which indicated whether their response was “precise” (error <= 15°) or “imprecise” (error > 15°). Training data were not used for further analyses.

Following a brief break (typically 20 min), participants performed the main task, without behavioral feedback, but with concurrent EEG recordings (see next). Each participant completed 6 blocks of the task (100 trials each), with periodic breaks at the end of every set of 50 trials. Thus, in total, we acquired and analyzed 14,400 trials of behavioral data from 24 participants.

#### Eye-tracking and gaze-based trial exclusion

For all participants, gaze fixation position was monitored binocularly at 500 Hz, and stored offline for analysis. In *post hoc* analyses, any trial in which the subject’s gaze deviated by more than 1 dva from the fixation dot along the azimuthal direction was flagged, and excluded from further analysis. Given that we were primarily interested in the effect of distractors on visual working memory, this criterion for gaze-based exclusion was applied only in a window around (100 ms before to 400 ms after) distractor onset during the delay epoch. We did not reject trials based on gaze deviation during other times in the delay epoch, because, barring the fixation dot and placeholders, there were no other stimuli on screen during this epoch. We also did not reject trials based on gaze deviation during the memory array presentation, because there was no prior information (or pre-cue) that could have systematically biased gaze toward either grating. In addition, no gaze-based rejection was performed for trials in which the distractor did not appear. Finally, for 6/24 participants gaze fixation could not be reliably recorded, largely because these participants wore eyeglasses with strongly reflective lenses; trial rejection rates for these participants were >15%. To avoid major biases in the statistical distribution of task parameters, for these 6 participants, we included all trials in the analyses; the results were similar even with these participants excluded from the analyses. For the remaining participants (n=18/24), the median eye-tracking rejection rate was 5.2%±0.8% (mean±s.e.m.).

### Behavioral data analyses

#### Estimating precision and bias in behavior

Participants reported their best estimates of grating orientation. Simple behavioral metrics, computed directly from the data, revealed the effects of the distractor on precision and bias in participants’ reports^66–68^, without the need for fitting complex psychophysical models^69^.

We computed the effect of the distractor on the precision of each participant’s orientation report using the circular standard deviation (CSD)^68^, a circular measure of dispersion of the orientation reports about its true value. A smaller CSD indicates higher average precision in orientation reports. Data were pooled from trials across all blocks for this analysis. We computed and compared the CSD separately for the trials in which the distractor appeared on the same hemifield, versus the opposite hemifield, of the probed stimulus; this analysis was performed separately for the cued and uncued grating reports. In addition, we fit a mixture model comprising von Mises and uniform distributions to estimate precision from the pooled responses, separately for the distractor-same and distractor-opposite trials. The difference (Δκ) in or modulation index (MI-κ; defined in the Results) of precision between the distractor-same and distractor-opposite trials was quantified separately for the cued and the uncued memoranda, and compared with permutation tests (see Methods section on *Statistical Tests*).

We also computed the effect of the distractor in biasing orientation reports for the probed grating, following a standard protocol^28,67^. First, we plotted the median orientation report error for the probed grating (reported – actual probed orientation) as a function of the angle difference between the distractor and the probed grating (distractor – actual probed orientation); this computation was performed with trials pooled in overlapping bins (size: ±15°, shift: 1°; range: −90° to 90°), based on the difference between the distractor and the probed grating orientation. Note that, in Figure 1E, if orientation reports were systematically biased towards the distractor orientation, then the y-axis (orientation report error) would be positive for positive values of the x-axis (relative distractor orientation), and vice versa for bias away from the distractor orientation. We quantified this bias by computing the signed area under the curve, with a procedure similar to that employed in previous studies (e.g., Myers et al^28^). Briefly, the area under the curve in quadrants II and IV were summed and subtracted from the summed area under the curve in quadrants I and III (Fig. 1E) – relative orientation values from −10° to 10° were excluded in this analysis because the distractor’s orientation was at least 10° away from either of the memory array grating orientations (see Methods, Behavioral Data Acquisition section). This analysis was performed both with data from all trials, and also separately for trials in which cued and uncued gratings were respectively probed. For visualization purposes, the bias curves were fit with a Gaussian derivative function^67^.

#### Estimating the contribution of distractor orientation reports

To quantify the contribution of distractor orientations reports to the observed behavioral responses, we fit, for each participant, a three-component mixture model – comprising two von Mises distributions and a uniform distribution ^29^:

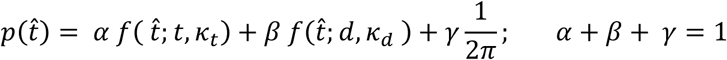

*t̂* denotes the reported orientation, *p*(*t̂*) the density (fit) of orientation reports, *f*(*t̂*; *t*, ***κ***_*t*_) is a von Mises distribution centered on the target orientation *t*, with concentration parameter ***κ***_*t*_, *f*(*t̂*; *d*, ***κ***_*d*_) is a von Mises distribution centered on the distractor orientation *d*, with concentration parameter ***κ***_*d*_, the uniform distribution models random guesses, and (*⍺*, *β*, *γ*) represent, respectively, the mixture weights associated with the target distribution, distractor distribution, and the uniform weight; these are constrained to sum to 1. We estimated mixture weights and concentration parameters (***κ***_*t*_, ***κ***_*d*_) using expectation–maximization^29^ for the response distribution associated with the cued memorandum, and separately for the distractor-same and distractor-opposite trials. We call this the “target + distractor” model; the “target” refers to the cued memorandum, when it was probed for recall.

Next, to evaluate the contribution of distractor orientation reports, we fit a simpler mixture model excluding the distractor von Mises component; we term this the “target alone” model:

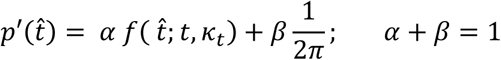

The two models – “target+distractor” and “target alone” were compared using the corrected Akaike Information Criterion (AICc), as well as the Bayesian Information Criterion (BIC) ^31,32^. Briefly, the two metrics trade off goodness-of-fit – quantified with the log-likelihood – against model complexity – based on the number of model parameters – with BIC imposing a more stringent penalty on model complexity. AICc and BIC values were computed separately for the distractor-same and distractor-opposite conditions, and then summed. Finally, we also repeated these same model comparison analyses by fitting both “target+distractor” and “target alone” models omitting the uniform component, and obtained results similar to those with this term included (see Results).

### EEG data acquisition and preprocessing

#### EEG acquisition

High density EEG data was acquired with the Active Two recording system (Biosemi, Amsterdam, Netherlands), with 128 Ag-AgCl active electrodes (Biosemi Inc.). During recording, the data were referenced online to the Common Mode Sense (CMS) active electrode. The data was recorded continuously throughout the task with a sampling rate of 4096 Hz and stored for offline analysis. EEG data for one participant could not be retrieved due to a critical hardware error. EEG analyses were performed with the remaining 23/24 participants.

#### EEG Preprocessing

EEG data was preprocessed in MATLAB using the EEGLAB^70^, ERPLAB^71^ and Noise Tools (http://audition.ens.fr/adc/NoiseTools/) toolboxes. The data was first resampled at 256 Hz (*pop_resample* function, EEGLAB), bandpass filtered from 0.05 to 40 Hz using Butterworth filter of order-2 (*pop_basicfilter*, ERPLAB), and cleaned of line noise at 50 Hz (*pop_cleanline*, EEGLAB). Next, noisy electrodes were identified as those having more than 33% of their values >4 times the median absolute value across all electrodes and time points (*nt_find_bad_channels,* Noise Tools); this yielded a channel removal rate of 4.1% ± 0.8% [mean ± s.e.m.] on average, per participant. The removed electrodes were then interpolated using the neighboring electrodes’ data with spherical spline interpolation (*pop_interp,* EEGLAB). Subsequently, the trials were epoched from −1600 to 2300 ms relative to the cue onset and median re-referenced by subtracting the median value across all electrodes^76^. The epoched data were then baseline normalized by subtracting the mean activity from −200 to 0 ms relative to cue onset from all time points, separately for all electrodes and trials (*pop_rmbase,* EEGLAB). Next, we applied independent component analysis (ICA) using the infomax algorithm (*pop_runica*, EEGLAB) to identify artifacts arising from physiological and other, non-neural sources. We used ICLabel^72^, an EEGLAB extension that probabilistically identifies ICs corresponding to neural sources versus those arising from eye movements, muscle artifacts, channel noise and other noise sources. Any component belonging to a non-neural source with high probability (>70%) was flagged, and removed after visual inspection; 3.2 ± 0.6 [mean ± s.e.m.] components were removed on average, per participant. In a final pass, we identified noisy trials using the SCADS algorithm^73^.

Briefly, SCADS identifies outlier signal values based on statistical measures like the standard deviation, maximum amplitude, and the like. Trials in which SCADS flagged >20 percent of electrodes (outlier criterion, λ=5.0) were removed from the analysis. In other trials, noisy electrodes were linearly interpolated based on the nearest 4 neighbors; this yielded a median trial removal rate of ∼3% trials per participant.

### EEG data analyses

#### Decoding stimulus orientation

We decoded the orientation of memory array and distractor gratings based on multivariate pattern analysis of EEG responses. Whereas the memory array grating orientations were decoded both during memory array presentation (Fig. 1A) and the delay epoch (Fig. 1A), distractor grating’s orientation was decoded only during its presentation epoch (Fig. 1A). The orientations of the memoranda were decoded with 14 posterior electrodes over the hemisphere contralateral to the respective stimulus (Fig. 2B, inset) pooled across the cueing conditions (see also control analysis, next, in which occipital electrodes were used bilaterally for decoding). The distractor’s orientation was decoded with all^39^ posterior electrodes over occipital cortex (e.g., Fig. 3A-B). We employed a multivariate decoder based on the Mahalanobis-distance, following a procedure established in literature^12,29,30^. Briefly, EEG data were downsampled to 128 Hz by taking an average of each set of two successive samples (original sampling rate: 256 Hz). To improve the decoding accuracy, we used a sliding window (size: 300 ms, shift: 8 ms) and concatenated all the data points in that time window from all electrodes into a one-dimensional vector (38 time points in each time window x number of electrodes). We utilized the pattern of temporal variability of the EEG response within each time window for decoding^30.^ Therefore, we subtracted the mean activity across the time window for each channel, and applied principal component analysis (PCA) to reduce the number of feature dimensions; the top-k PCA dimensions that explained 95% of the variance across trials were retained for decoding (“encoding vector”).

For training and evaluation, we employed a leave-one-trial-out cross-validation approach. For each participant, the model was trained on all but one trial, with orientations binned relative to the orientation in the left-out trial into one of 16 different bins (bin centers: 11.25° apart, bin width: ±30°). Then the Mahalanobis distance was computed between the test trial’s encoding vector and the mean encoding vector of each training bin. The computed distances were de-meaned and the sign of the distances was reversed for ease of visualization. To obtain a smooth decoding estimate, this procedure was repeated 8 times by sliding the bin centers by 1.4° each time, and the resultant decoding profiles were averaged across repetitions (e.g., Fig. 2A). To estimate decoding accuracy, the sign-reversed distances were multiplied elementwise by a cosine kernel, symmetric around 0° (period: 180°), and averaged. This procedure was performed for each participant, memory array location (left/right), distractor location (left/right), and each time window, separately.

#### Distractor-evoked ERP estimation

The event related potential (ERP) was measured by signal-averaging across the same set of 14 contralateral (or ipsilateral) electrodes as employed in the decoding analyses. First, the data were high-pass filtered with a lower cut-off at 0.5 Hz using an FIR filter (N=500). Subsequently, the data were epoched from 800 ms before to 700 ms after distractor onset and then averaged across channels and trials to obtain the ERP waveform. This was done separately for each of the four distractor conditions (cued/uncued x left/right hemifield). Click or tap here to enter text. The N1 amplitude was defined as the mean signal value in a window extending from 150 to 200 ms following distractor onset and P2/P3a amplitude as the mean signal value in a ±40 ms window around the respective P2 and P3a peaks ^31,33^.

#### Median-split analyses based on memorandum maintenance, distractor encoding strength, timing and gating

We tested the association of diverse neural metrics and task variables to distractor-induced behavioral bias. For example, we measured the association between the strength of the cued memorandum’s maintenance and behavioral bias. For this analysis, the strength of the cued memorandum’s maintenance was quantified based on the average decoding accuracy (cosine similarity) in a 500-ms window in the delay period – beginning with cue onset, until 100 ms preceding the earliest distractor onset. Then, we divided trials into “weak” and “strong” memorandum maintenance trials based on a median split of the cosine similarity metric; the median split was performed for each participant and for each distractor condition (cued/uncued x left/right hemifield) separately. Weak and strong memorandum maintenance trials were then aggregated across the conditions, and the behavioral bias in the cued orientation report was computed separately for the two groups of trials with a procedure identical to that described previously (see Methods section on *Estimating precision and bias in behavior*).

The association between other metrics like the cued memorandum encoding strength, uncued memorandum maintenance strength, distractor encoding strength, distractor timing or the strength of gating and behavioral bias was computed in an identical manner, except that the median split was performed based on the respective metric. For the cued memorandum encoding strength, a median split was performed based on cued memorandum decoding accuracy values in a window from 100 ms to 500 ms following memory array onset. For the uncued memorandum strength, a median split was performed based on uncued memorandum decoding accuracy values in a 500 ms delay-period window, identical to that for the cued memorandum. For distractor encoding strength, a median split was performed based on distractor decoding accuracy (cosine similarity) values computed in a window from 100-400 ms after distractor onset. For distractor timing (early versus late), a median split was performed based on the times of distractor onset relative to the cue. For distractor input gating, the median split was based on the amplitudes of the distractor-evoked P2/P3a component over the electrodes contralateral to the distractor. Statistical tests were performed both to assess whether the bias for each half of the median split was, individually, significantly different from zero, and also to assess for within-participant (pairwise) differences across the median split halves. For each split, we computed one-tailed Bayes Factors (see next), which enabled us to quantitatively evaluate evidence for or against the effect of each factor on distractor-induced bias.

Prior to this analysis, we first tested and confirmed that these factors were independent of each other and of the distractor’s encoding strength. First, trials were divided into high and low cued memorandum maintenance groups based on a median split, and we tested whether distractor encoding strength differed significantly between these two groups. Second, trials were split based on the timing of distractor appearance (early vs. late), and we assessed whether this timing influenced distractor encoding strength or distractor gating. Finally, trials were divided based on distractor encoding strength (strong vs. weak), and we examined whether distractor gating strength differed between these groups. These control analyses confirmed that the primary factors of interest were not confounded with one another, providing a basis for interpreting subsequent analyses.

### Computational model of distractor biases in WM

#### i) Network architecture

We synthesized our behavioral and neural findings by modifying and extending a bump attractor model of working memory^12^. The visual cortex (VC) and higher cortex (HC), for each of the cued and uncued visual hemifields, were modeled with distinct ring attractors (total of 4 ring attractor models). Each attractor model comprised 1024 neurons in total out of which 512 were excitatory (E) and 512 were inhibitory (I). Each neuron was described by a standard rate model and represented stimulus orientation topographically, depending on its location along the ring (Fig. 5 A, top; θ ranging from −90° to +90°).

For ease of illustration, we start by describing the dynamical equation governing the firing rate r of the cued visual cortex attractor (VC,C) as a function of time *t*^27,28^.

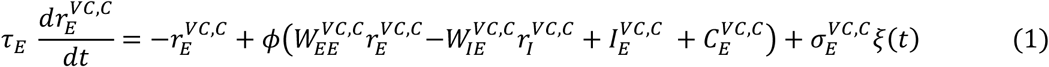

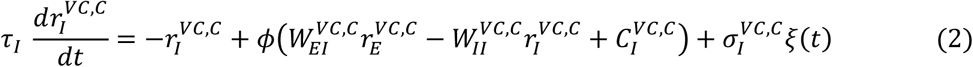

Where, 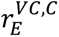 and 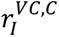 denote the firing rate of the excitatory and inhibitory neurons of the visual cortex, cued attractor (VC,C) respectively. Τ_*E*_ and τ_*I*_ are neuronal (integration) time constants for E and I neurons respectively, 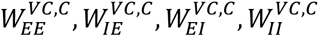 are the connection strengths from E to E, I to E, E to I and I to I neurons respectively for the VC,C attractor. 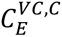 and 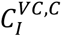 are the constant input bias currents to E and I neurons respectively of VC,C attractor. Ξ(*t*) represents Gaussian white noise with unit variance, scaled by parameters 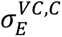 and 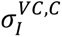 for E and I neurons of VC,C attractor respectively. 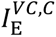 represents the input from other sources (explained in equations 3 and 4, see next). *ϕ* represents the function that maps the input current to the output firing rate adopted from Wimmer et al., 2014^27^

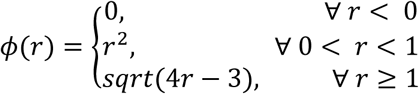

Three of the four sets of recurrent connections – *W*_*IE*_, *W*_*EI*_, *W*_*II*_ – were uniform and all-to-all (i.e., *w*^*ij*^ = *w*_0_ ∀ *i*, *j*). On the other hand, recurrent excitatory connections 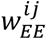 were local, and followed a von Mises profile: *exp*(***κ**cos*(*θ*_*i*_ − *θ*_*j*_)) where ***κ*** is the concentration parameter and *θ*_*i*_ and *θ*_*j*_ are the orientation preference of excitatory neurons *i* and *j* respectively.

The firing rates of three other attractors, namely, the visual cortex uncued attractor (VC,U), higher cortex cued attractor (HC,C) and higher cortex uncued attractor (HC,U) also follow identical dynamical equations. The values of the parameters for the 4 different attractor models are shown in SI Table S1.

Inputs to the cued visual cortex attractor (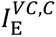, equation 1) comprised 3 contributions:

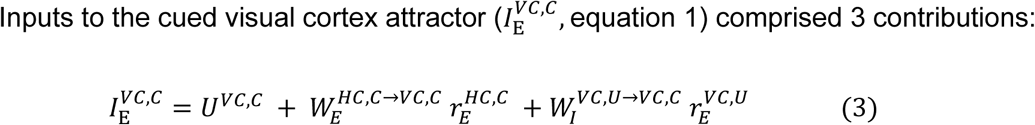

The first term, *U*^*VC*,*C*^ represents the bottom-up (feedforward) inputs due to the external stimulus that become active during the presence of the memory array and distractor gratings (successively). The second term, 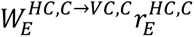 represents top-down (feedback) inputs from the HC to the VC attractor, organized topographically (one-to-one); here, 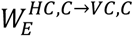 denotes the top-down excitatory connection weights from the HC,C to VC,C attractor. The third term, 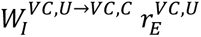 represents long-range, lateral inhibitory inputs received from the VC attractor in the opposite hemisphere (cross-hemispheric inhibition); here, 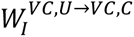 represents the lateral inhibitory connection weights from the VC,U to VC,C attractor. Note that the magnitude this inhibition modulates with the firing rate of the excitatory neurons (*r*_*E*_) in the opposite hemifield; an intervening inhibitory population that converts this long-range excitatory projection into functional inhibition (via weights, *W*_*I*_) was not explicitly modeled. The inputs 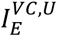 to the uncued visual cortex attractor also followed an identical equation except for minor modifications to the respective bottom-up inputs (*U*^*VC*,*U*^), top-down inputs 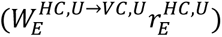 and long-range, lateral inhibitory inputs 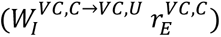.

Inputs to the cued higher cortex attractor (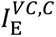, equation 1) comprised only one contribution corresponding to bottom-up connections from the VC,C attractor, also organized topographically (one-to-one):

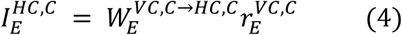

Where 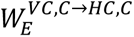 represents the topographic bottom-up connection strength from the E neurons of the VC,C to the E neurons of the HC,C attractor 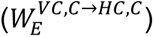. The inputs 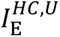 to the uncued higher cortex attractor also followed an identical equation except for its bottom-up inputs 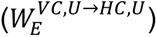.

#### ii) Modeling the distinct task epochs

##### Encoding

500 ms after the trial onset the memorandum was presented for a duration of 500 ms. The visual cortex (VC) attractor receives sensory input during this period (through *U*^*VC*,*C*^ and *U*^*VC*,*U*^terms in equation 3). We modeled this by changing the spatial profile of the input current to the excitatory neurons of the VC attractor according to:

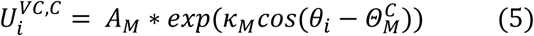

Where, 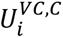 is the input current to the excitatory neuron *I* of VC,C attractor, *A*_*M*_ is the amplitude of the input current, ***κ***_*M*_ is the concentration parameter determining the spread, *θ*_*i*_ is the orientation preference of excitatory neuron *i* and 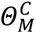 is the memory array grating’s orientation on the cued hemifield. The spatial profile of the input current was centered on the E neuron representing the respective grating’s orientation. This instantiated localized bumps of activity in VC, at locations corresponding to the orientations of the encoded gratings. A similar equation was used to model the input current for the VC,U attractor. The PFC attractor did not receive any direct inputs other than the bottom-up input received from the VC attractor during encoding (equation 4).

##### Delay

The delay period commenced immediately after memory array offset. During the delay period, until the end of the trial, we disconnected the bottom-up connections from the VC to the HC attractor (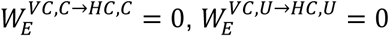 in equation 4), rendering the HC attractor immune to subsequent sensory input, thereby modeling “strong” distractor gating. These connection values were set to 0.01 only for the specific case of modeling “weak” distractor gating (Fig. 6E, SI Fig. S5F). 600 ms after the memory array onset, the cue appeared. To mimic prioritization of cued information in WM^24^, we reduced the noise terms for both the cued attractors (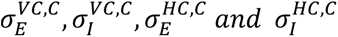in equations 1 and 2) by 60% and increased top-down feedback to cued VC attractor (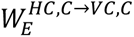 in equation 4) by 20%, following cue onset.

##### Distractor presentation

The distractor was presented for a duration of 0.1 seconds, at a random time (distributed exponentially) between 1200 ms and 1950 ms after the memory array onset, mimicking the behavioral paradigm. The distractor was presented either in the cued or the uncued hemifield with equal probability and was encoded by the visual cortex (VC) attractor representing the corresponding visual hemifield. Distractor presentation was modeled by adjusting the spatial profile of the input current to the excitatory neurons of the VC attractor as follows:

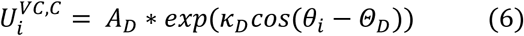

Where 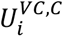 is the input current to the excitatory neuron *i* of VC,C attractor, *A*_*D*_ is the amplitude of the input current, ***κ***_*D*_ is the concentration parameter determining the spread, *θ*_*i*_ is the orientation preference of excitatory neuron *i* and *Θ_D_* is the distractor’s orientation. In addition, we enhanced the cross-hemifield inhibitory weights (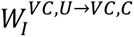 or 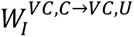 in equation 3) multiplicatively (by ∼30x) during distractor presentation; this enabled modeling the bias away from the distractor’s orientation observed in the distractor-opposite condition.

#### iii) Quantifying simulated behavioral and neural metrics

We ran a total of 10,000 simulations with a different random seed each time; all results represent an average of these simulations. During each simulation, we randomly sampled the memory array (cued and uncued) and distractor orientation from a circular uniform random distribution. The “neural” orientation maintained by the network was decoded using the population vector representation in the visual cortex attractor given by:

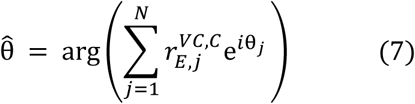

where, θ̂ is the decoded orientation from the VC,C attractor. *N* is the number of E neurons. 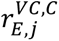 and θ_*j*_ is the firing rate and preferred orientation respectively of neuron *j* in the VC,C attractor. Orientations from uncued visual cortex attractor were also decoded in an identical manner, using its respective population vector. The model’s neural orientation decoding was computed with the VC attractors alone to mimic the decoding of neural orientation from only the occipital electrodes in the EEG data. By contrast, drawing inspiration from recent work that has highlighted the involvement of both sensory (visual) cortex (VC) and higher brain areas (HC) in robust working memory maintenance^3–5,9,13,18,41^, the decoded orientation from the VC and HC attractors at the end of the delay period were averaged to generate the model’s behavioral orientation estimate (see SI Results Section); orientation from the higher cortex attractor were decoded in a manner identical to that in equation 7, using their respective population vectors.

### Statistical Analyses

We employed a non-parametric Wilcoxon signed rank test for comparing median errors in orientation reports (Fig. 1B, C). Permutation tests (10000 shuffles, unless otherwise indicated) were performed for behavioral bias across conditions; one-sided p-values are reported under the *apriori* hypothesis that the attractive (repulsive) biases occur when the distractor appeared on the same (opposite) hemifield as the memorandum (Fig. 1G-H; 2C-D; 3C-D, 4B-C). All the other tests were two-sided, unless otherwise stated. To study the effect of distractor condition and cueing on precision in visual WM (Fig. 1C), we performed a 2-way ANOVA with the circular standard deviation as the dependent variable and distractor condition (distractor-same, distractor-opposite and no-distractor) and cueing (cued and uncued) as predictors; participants was treated as random effects. *Post-hoc* analyses were performed by averaging the values across the cue conditions and comparing CSD values for distractor-same versus distractor-opposite conditions with Wilcoxon signed rank test. Permutation tests by shuffling (1000 shuffles) distractor-same and distractor-opposite labels, were used to compare precision degradation induced by the distractor across cued and uncued memoranda. To find significant time points in decoding accuracies plotted as a function of time (Fig. 2B), we used the two-sided cluster-based permutation test^79^ with a cluster-forming threshold of p<0.05 and with 10,000 permutations of the data. The distribution of decoding accuracies across cued and uncued conditions was compared using permutation tests (two-sided) both for memorandum decoding and distractor decoding analysis (Fig. 2B, 3B). The distractor related ERPs across cued and uncued conditions were compared using the signed-rank test (Fig. 4A).

In addition, we also report the Bayes factor (BF^80^), a statistical measure that compares the likelihood of the alternate hypothesis to that of the null hypothesis under the observed data. BFs were computed with a JZS prior, using the Bayes Factor toolbox^81^. One-tailed BFs were computed specifically when comparing the effect of distractor features on behavioral biases; other BFs were two-tailed, unless otherwise stated. Moreover, one-tailed BFs, and associated p-values, were also used to compare the strength of biases across the median splits, because we had clear, *apriori* hypotheses regarding the effect of each memorandum (decodability) or distractor characteristic (timing, decodability, gating) on bias magnitude. BFs computed based on a two-tailed versus one-tailed tests are reported as BF_10_ versus BF_+0_/BF_-0_, respectively; the subscripts +0 and −0 denote the directionality of the hypothesis tested (greater or lesser than zero, respectively).

## Data and code availability

Data and custom Matlab code to replicate the experimental findings as well as the computational modeling results are available at the following link: https://osf.io/dqr7m/?view_only=71feeb114f744f8c987d6a22f230905f

## Declaration of generative AI and AI-assisted technologies in the writing process

During the preparation of this work the second author used ChatGPT and Open AI as a language editing tool. After using this tool/service, the second author reviewed and edited the content as needed and takes full responsibility for the content of the parts of the publication that was co-written by them.

## Acknowledgements

We thank Rishabh Bajpai and Mirudhula Mukundan for their help in piloting the task design. This research was supported by a Wellcome Trust-Department of Biotechnology India Alliance Intermediate fellowship, DST Swarna-Jayanti fellowship, a Pratiksha Trust Intramural grant, an India-Trento Programme for Advanced Research (ITPAR) grant and a Department of Biotechnology-Indian Institute of Science Partnership Program grant (all to DS).

## Author contributions

D.S. and S.G. designed the study; S.G. conducted the experiment; D.R and S.G. analyzed the data and performed model simulations; D.R, S.G. and D.S. wrote the paper.

## Supplementary Information: Results

### Error-correcting dynamics in a two-tier attractor model

We present here additional simulations clarifying the two-tier architecture for the WM attractor model employed in our simulations. We employed a two-tier model, rather than a single tier model for modelling robust mnemonic maintenance. Recent literature suggests that mnemonic representations in higher order brain regions are less susceptible to interference from distractors than those in early sensory cortices^1–4^. Top-down feedback from the higher order brain region has been hypothesized to help mitigate distractor interference in the early sensory cortices^1,2,4–6^. To test whether a similar mechanism could also be modeled, we performed simulations (n=10^4^, across 10 runs) by varying the strength of top-down feedback from the HC to VC attractors in both hemifields (*W*^*HC*→*VC*^ term in equation 4, main text) by a factor of 2x. Indeed, increasing the top-down feedback to the VC markedly reduced maintenance error for both the cued and the uncued VC attractor (SI Fig. S6A, CSD reduced (%): VC,C=5.94±0.55%, range: 4.19 to 8.11%; VC,U= 8.47±0.68%, range: 4.61 to 12.12%). In our model, such error correcting dynamics from the HC were essential to overcome the disruptive effect of the distractor on the VC; they helped reinstate an accurate mnemonic representation in the VC attractor following distractor disappearance (Fig. 5C-D, main text).

Moreover, during recall, the behavioral feature estimate of the memorandum was computed from both VC and HC attractor representations. Recent research indicates that visual WM maintenance of involves multiple brain regions, including both sensory and higher-order areas^7–11^. It is possible, then, that a (weighted) average of the memorandum’s feature representation across multiple brain regions contributes to a more stable behavioral readout. We hypothesize that such a readout improves precision by cancelling out independent noise and drift in mnemonic representations across different brain regions. We tested this hypothesis by observing the average precision of “behavioral” readouts over n=10,000 simulations based on the decoded representations of each memorandum’s orientation, at the end of the delay period. In a first set of simulations, these readouts were estimated from the VC attractor alone (equation 7, main text). In a second set of simulations, we averaged the readouts from both the VC and the HC. The averaged readout was numerically more accurate as compared to the estimate from any single brain region alone (SI Fig. S6B, CSD reduction (%): Cued=1.18±0.2%, range: 0.58 to 2.68%; Uncued=2.12±0.15%, range: 1.48 to 2.71%). Consequently, in the simulations in the main text, we averaged the HC and VC attractor readouts to obtain the model’s behavioral readout.

## Supplementary Information: Figures and Tables

**Supplementary Figure S1.**
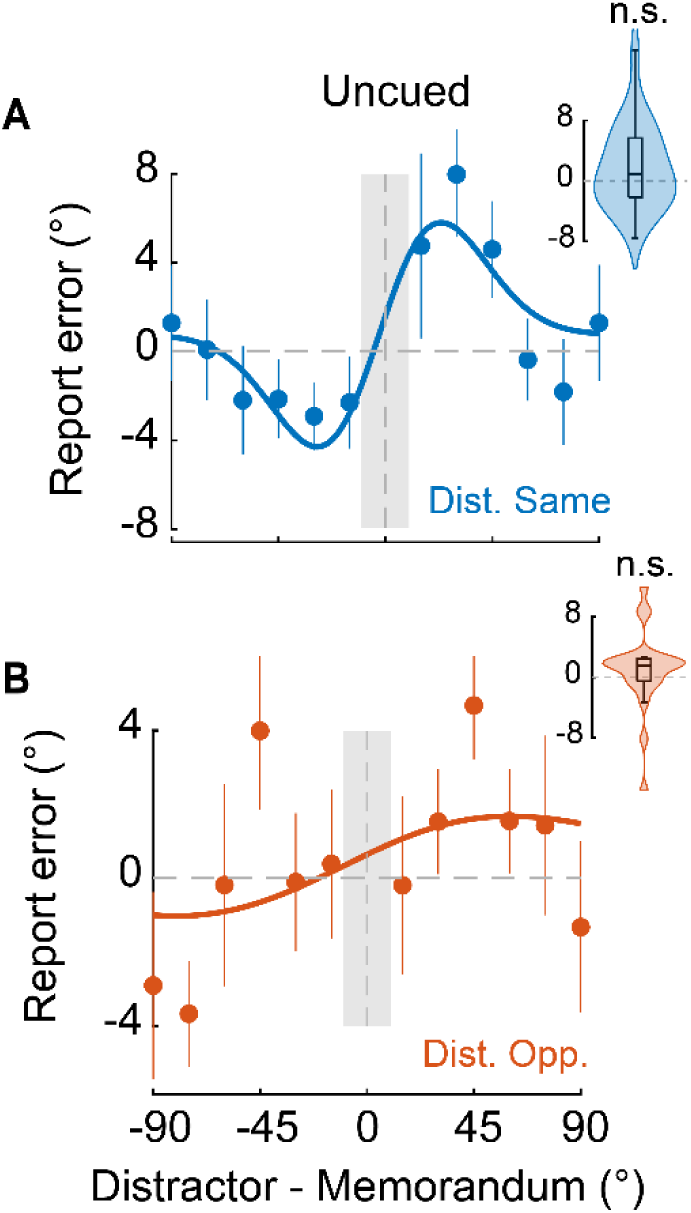
Distractor-induced behavioral biases for the uncued memorandum. **A.** Same as in Figure 1G (main text) but showing the average behavioral bias for the uncued memorandum when the distractor appeared in the same hemifield as the former. Other conventions are the same as in Figure 1G. **B.** Same as in panel A, but for trials in which the distractor appeared in the hemifield opposite to the uncued memorandum. Other conventions are the same as in panel A and Figure 1H (main text).

**Supplementary Figure S2.**
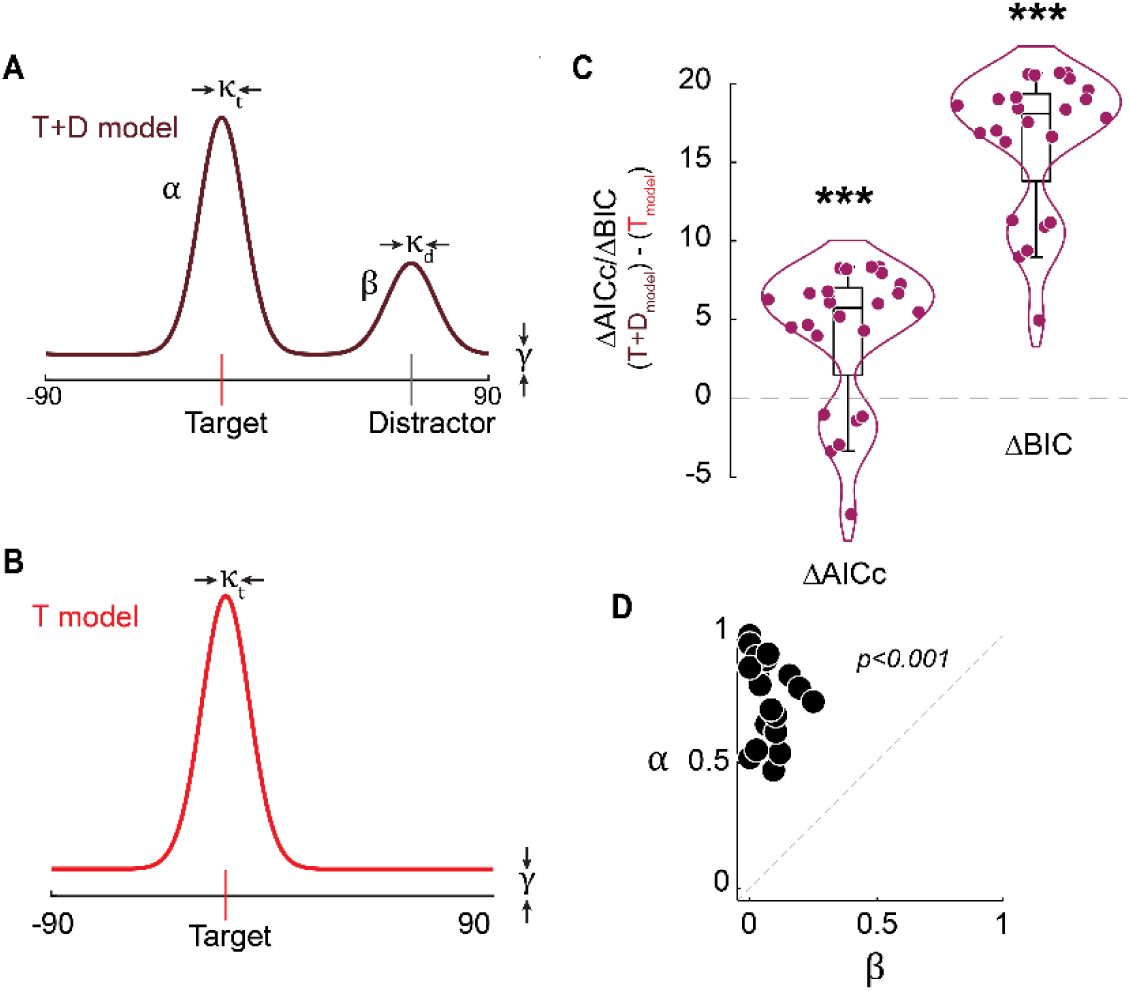
von Mises mixture model fit and evaluation. **A.** Schematic of a mixture model comprising two von Mises and a uniform distribution in which participants responses are modeled as a combination of target (memorandum) and distractor orientation reports, and guesses (“target+distractor”/T+D model). κ_t_ and κ_d_ denote the precision associated with the target and distractor responses respectively. *⍺* and *β* denote the contribution of the target and distractor to the orientation responses whereas, *γ* denotes the contribution of the uniform component (*⍺* + *β* + *γ* =1). **B.** Same as in A, but for a mixture model comprising one von Mises and a uniform distribution to model participants’ responses based on target orientation reports (“target alone”/T model) and guesses alone. Other conventions are the same as in panel A. **C.** Model comparisons. (*Left*) Difference between corrected Akaike information criterion (AICc) values for the “target+distractor” model fit and “target alone” model fit. A lower (more negative) value indicates a preferred model. Individual points: Participants. Violins, and box and whisker plots follow the same conventions as in Figure 1F-G (insets). (*Right*) Same as in the left panel but for the Bayesian information criterion (BIC). Asterisks: significant difference of median values between models, based on permutation tests. *p<0.05, **p<0.01, ***p<0.001, n.s.: not significant. **D.** Scatter showing the distribution of *⍺* (target weight) and *β* (distractor weight) obtained by fitting participants’ responses with the “target+distractor” combined mixture model. Points: Individual participants. Dashed diagonal line: line of equality.

**Supplementary Figure S3.**
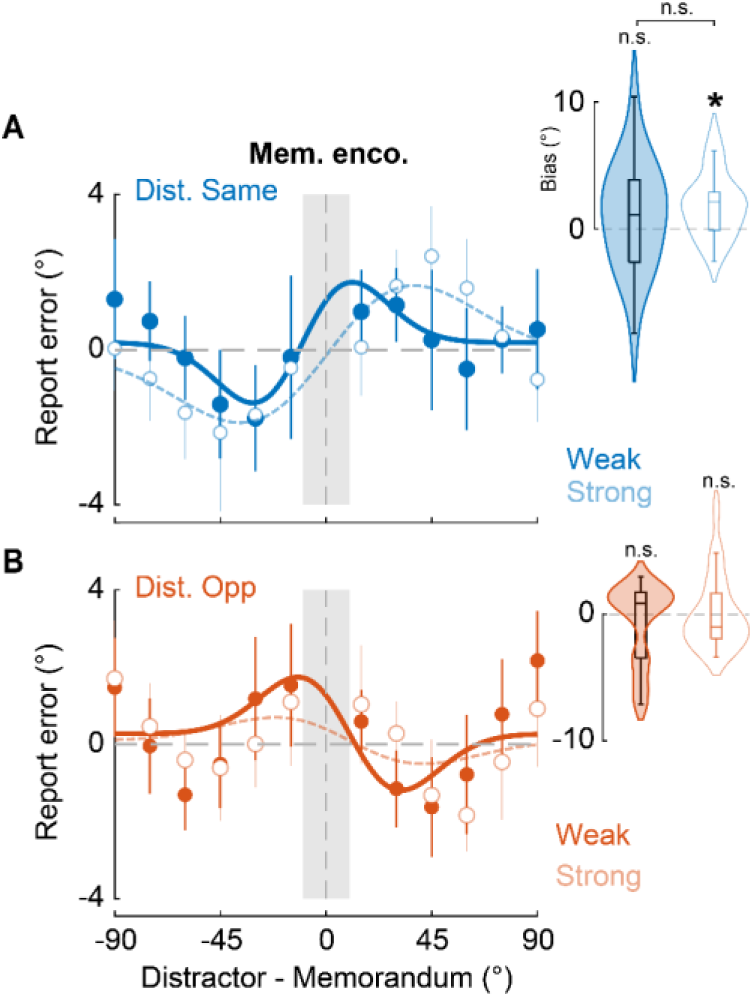
Effect of memorandum encoding on distractor-induced bias. **A.** Same as in Figure 2C (main text), but showing behavioral bias curves for the cued memorandum based on a median split of memorandum encoding strength – quantified with neural decoding accuracy following memorandum presentation (gray shaded bar, Fig. 2B, left) – when the distractor appeared in the same hemifield as the memorandum (n=23). Solid curve and filled violin plot (*inset*): Weaker memorandum encoding trials. Dashed curve and open violin plot (*inset*). Stronger memorandum encoding trials. **B.** Same as in panel A, but when the distractor appeared in the hemifield opposite to the cued memorandum. Other conventions are the same as in Figures 2C-D (main text).

**Supplementary Figure S4.**
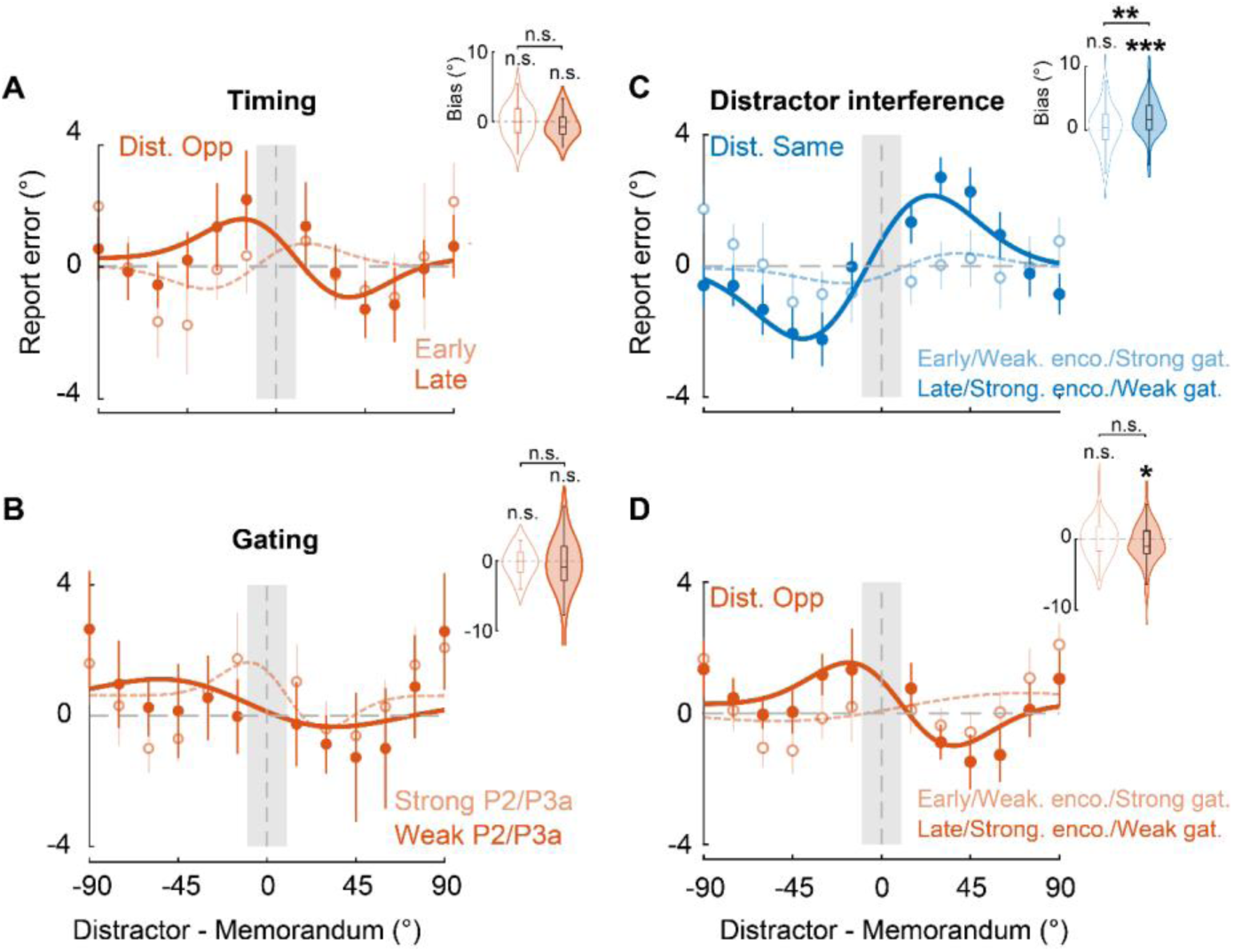
Distractor-opposite and omnibus analysis results. **A.** Same as in Figure 4B-C (main text) but showing the behavioral bias curves for the cued memorandum when the distractor appeared on the hemifield opposite to the cued memorandum – for trials in which the distractor appeared early (dashed, open symbols, unfilled violins) or late (solid, filled symbols, filled violins) during the delay period. **B.** Same as panel A, but for trials with strong (dashed) or weak (solid) distractor-evoked P2/P3a amplitude. (A-B) Other conventions are the same as in Figures 4B-C. **C.** Same as panel Figure 4B (main text), but showing the results of an omnibus analysis combining across conditions with high distractor interference – late timing, strong encoding, weak gating (solid, filled symbols, filled violins) – or low distractor interference – early timing, weak encoding, strong gating (dashed, open symbols, unfilled violins) – for trials in which the distractor appeared on the same hemifield as the cued memorandum. **D.** Same as panel C, but when the distractor appeared on hemifield opposite to the cued memorandum. (C-D), other conventions are the same as in Figure 4B (main text) and SI Figure S4A. (All panels) *: p<0.05; **: p<0.01; ***: p<0.001; n.s.: not significant.

**Supplementary Figure S5.**
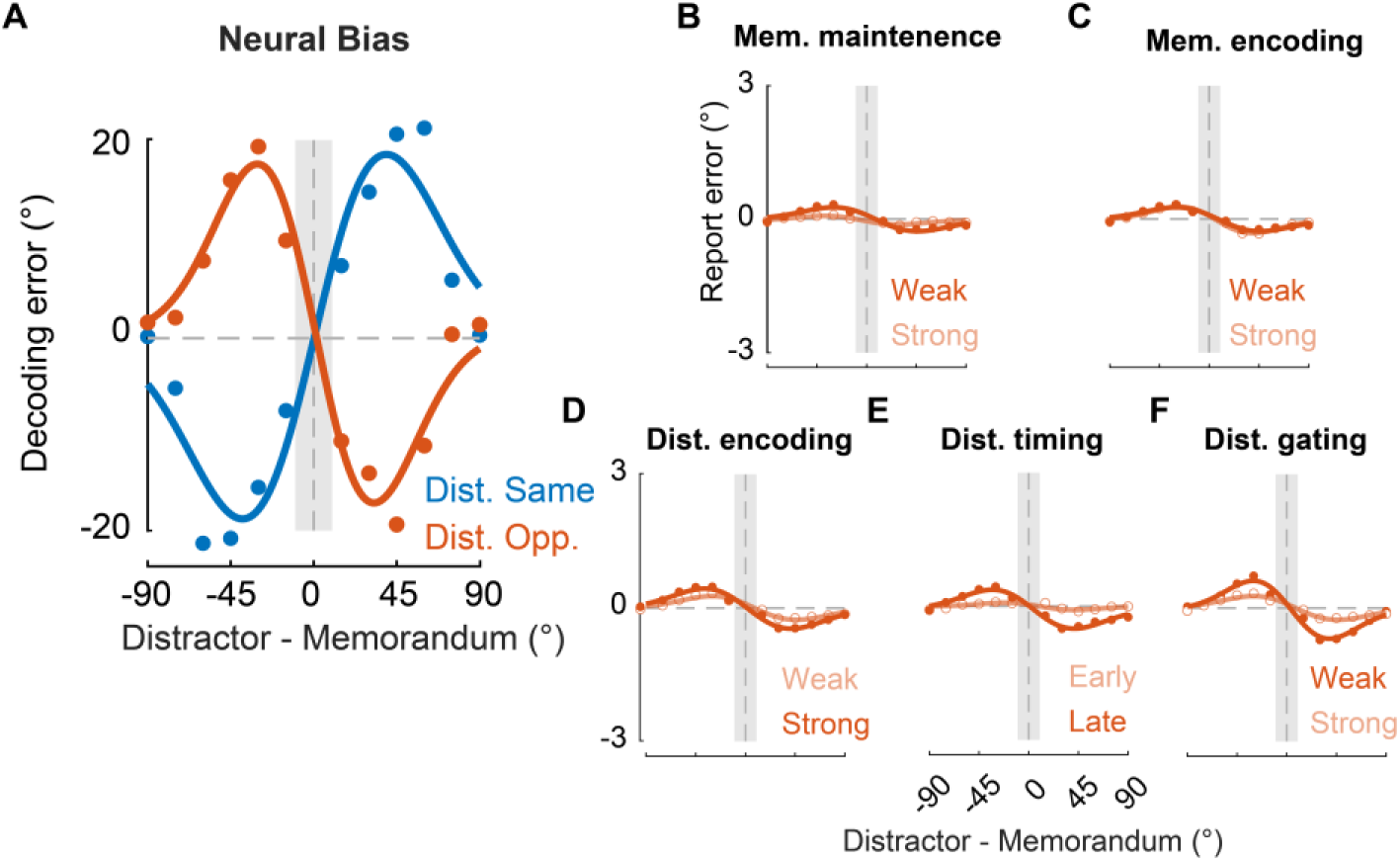
Simulations of neural bias and distractor-opposite effects. **A.** Simulated neural bias for the cued memorandum when the distractor appeared in the same hemifield (blue) or the opposite hemifield (orange) as the former. Other conventions are the same as in Figure 6A (main text). **B-F.** Same corresponding panels as in Figure 6B-F (main text), except showing simulated behavioral bias when the distractor appeared in hemifield opposite to the respective memorandum. Other conventions are the same as in corresponding panels in Figures 6B-F.

**Supplementary Figure S6.**
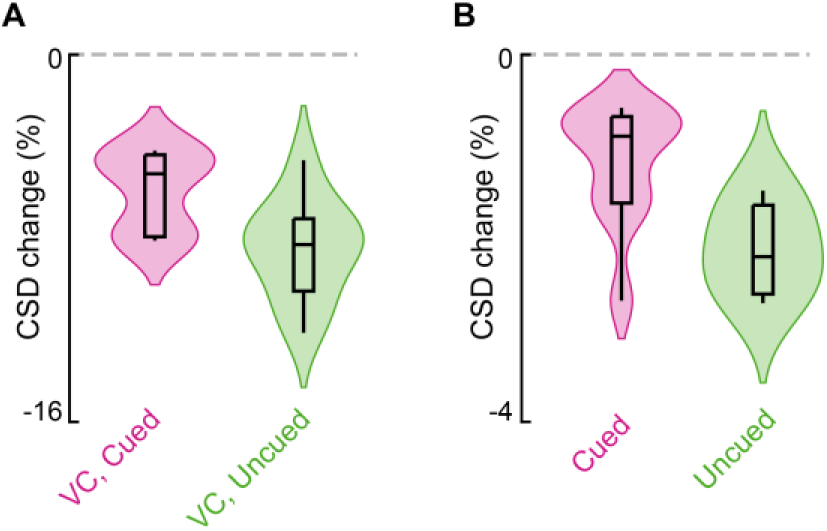
Error-correcting dynamics in WM. **A.** Change in error in the VC attractor readout upon incorporating top-down feedback from the HC attractor. Data shown separately for the cued (magenta) and uncued (green) attractors. Violin plots showing the distribution of values across 10 runs of the model (n=1000 simulations for each run); each simulation initialized with different random seeds, and values were averaged across simulations within a run. **B.** Change in error obtained by averaging readouts from the VC and the HC attractor relative to the mean readout error for either attractor (average of VC and HC readout errors). Other conventions are the same as in panel A.

**Supplementary Table S1.**
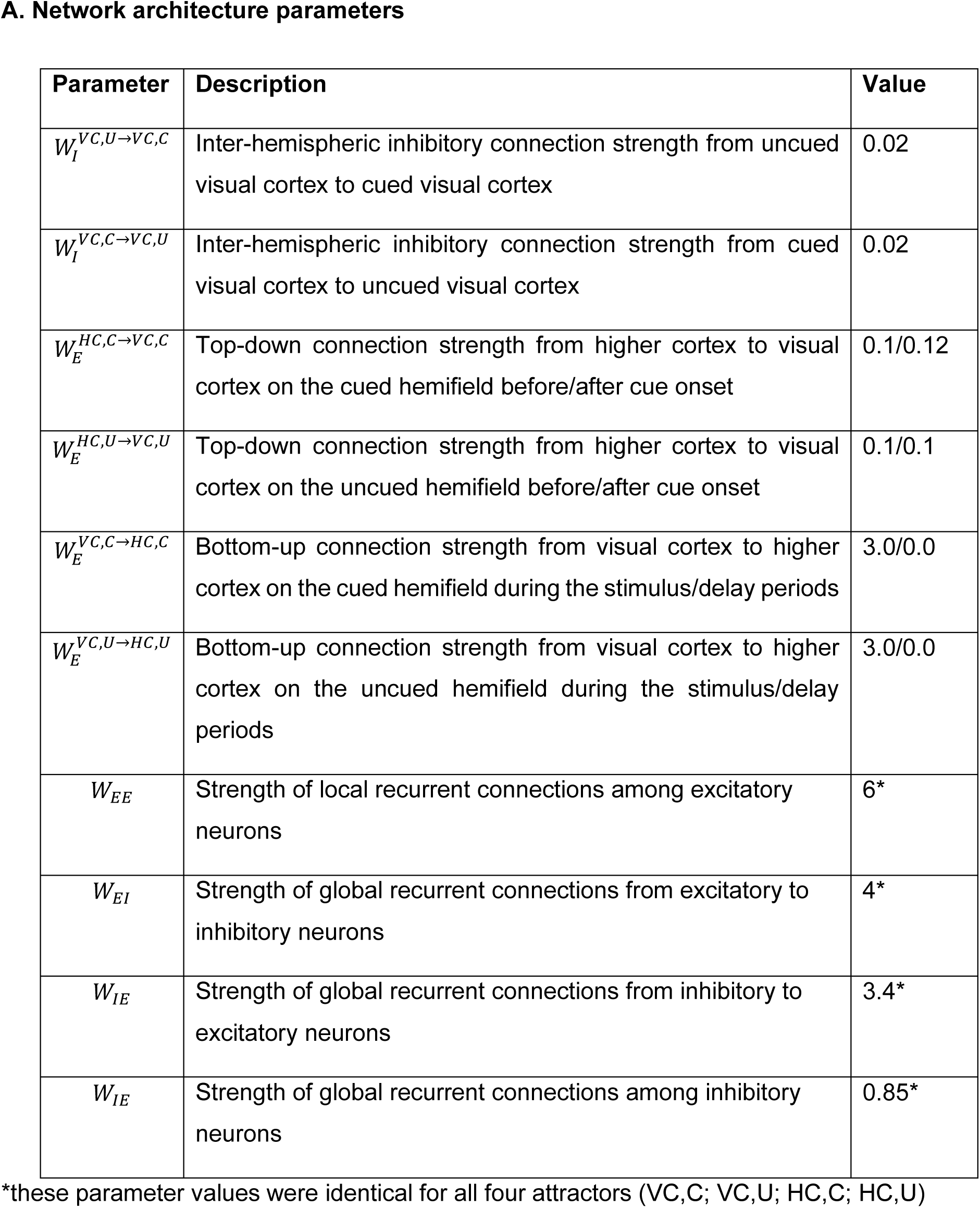
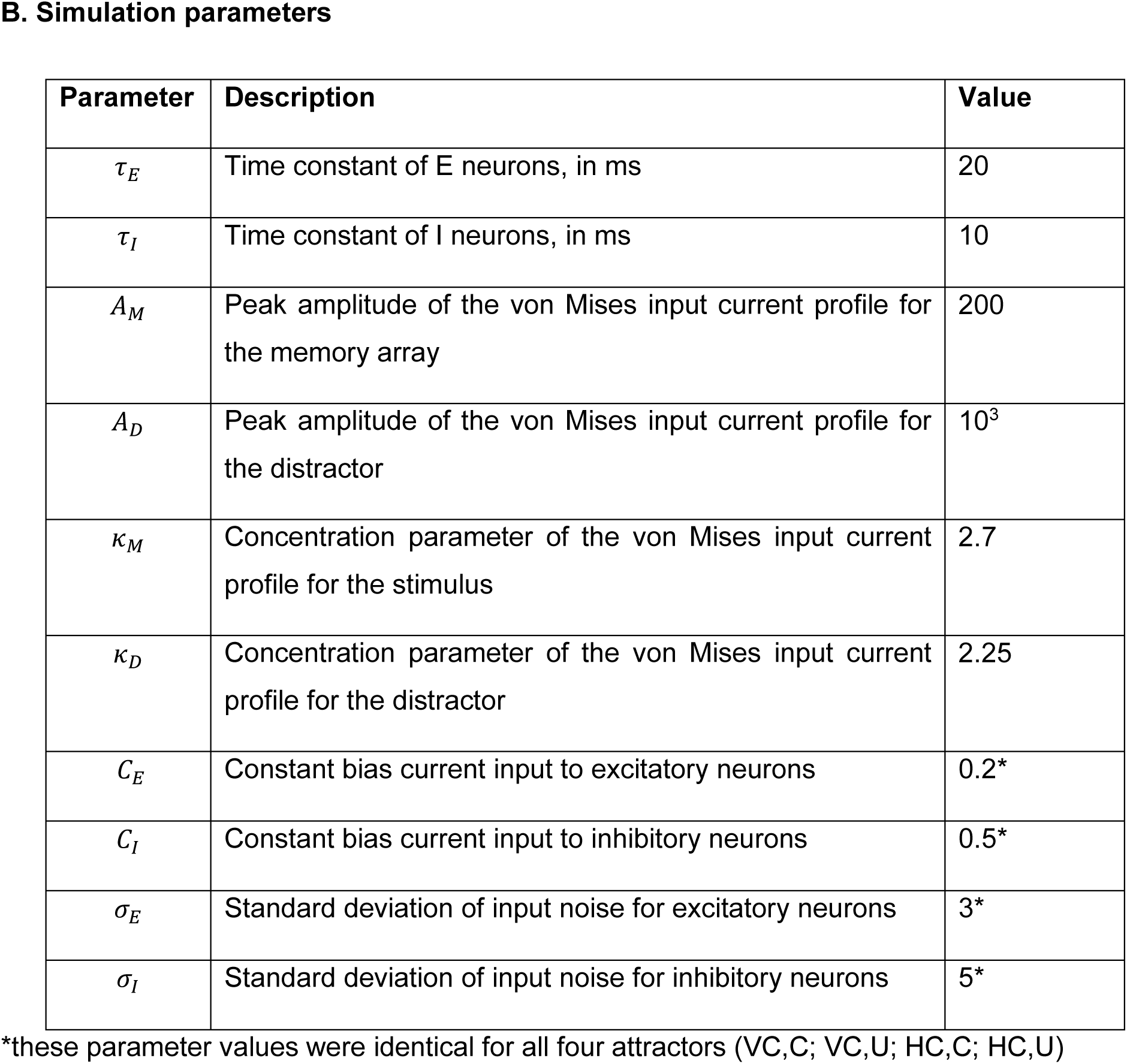
Network architecture and simulation parameters employed in the model.

## Notes

https://osf.io/dqr7m/?view_only=71feeb114f744f8c987d6a22f230905f

### Competing Interest Statement

The authors have declared no competing interest.

