## Supplementary Information for "Neural bases of space-specific distractor biases in visual working memory"

### Supplementary Information: Figures and Tables

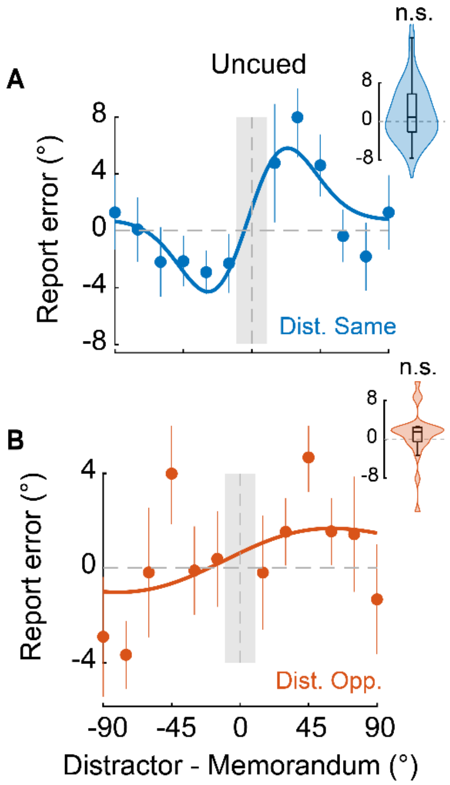


**Supplementary Figure S1. Distractor-induced behavioral biases for the uncued memorandum.**

**A.** Same as in Figure 1G (main text) but showing the average behavioral bias for the uncued memorandum when the distractor appeared in the same hemifield as the former. Other conventions are the same as in Figure 1G.

**B.** Same as in panel A, but for trials in which the distractor appeared in the hemifield opposite to the uncued memorandum. Other conventions are the same as in panel A and Figure 1H (main text).

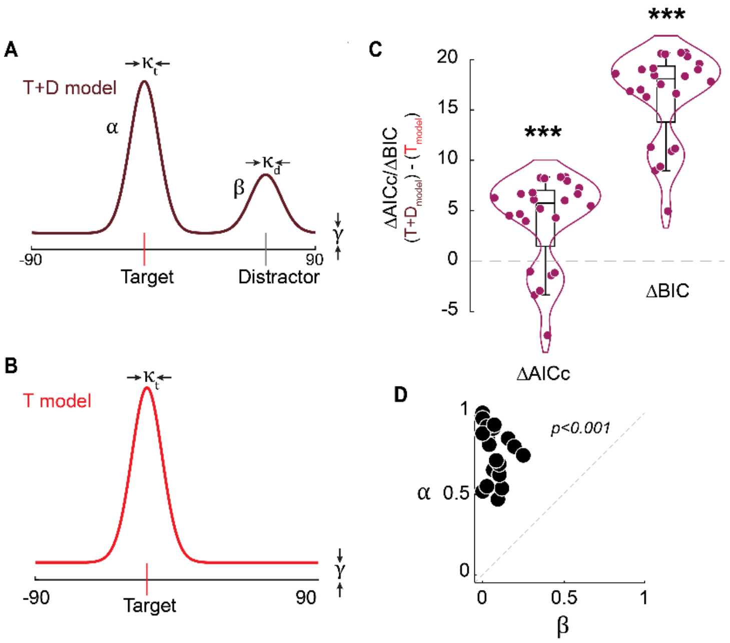


**Supplementary Figure S2. von Mises mixture model fit and evaluation.**

**A.** Schematic of a mixture model comprising two von Mises and a uniform distribution in which participants responses are modeled as a combination of target (memorandum) and distractor orientation reports, and guesses ("target+distractor"/T+D model). $\kappa_t$ and $\kappa_d$ denote the precision associated with the target and distractor responses respectively. $\alpha$ and $\beta$ denote the contribution of the target and distractor to the orientation responses whereas, $\gamma$ denotes the contribution of the uniform component ($\alpha + \beta + \gamma = 1$).

**B.** Same as in A, but for a mixture model comprising one von Mises and a uniform distribution to model participants' responses based on target orientation reports ("target alone"/T model) and guesses alone. Other conventions are the same as in panel A.

**C.** Model comparisons. (*Left*) Difference between corrected Akaike information criterion (AICc) values for the "target+distractor" model fit and "target alone" model fit. A lower (more negative) value indicates a preferred model. Individual points: Participants. Violins, and box and whisker plots follow the same conventions as in Figure 1F-G (insets). (*Right*) Same as in the left panel but for the Bayesian information criterion (BIC). Asterisks: significant difference of median values between models, based on permutation tests. *p<0.05, **p<0.01, ***p<0.001, n.s.: not significant.

**D.** Scatter showing the distribution of $\alpha$ (target weight) and $\beta$ (distractor weight) obtained by fitting participants' responses with the "target+distractor" combined mixture model. Points: Individual participants. Dashed diagonal line: line of equality.

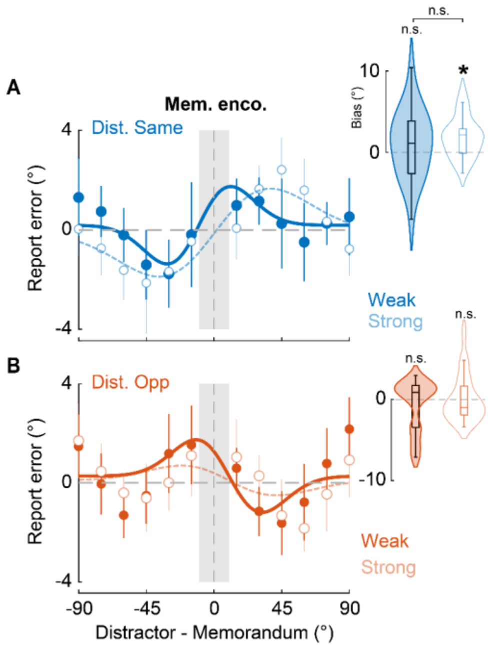


**Supplementary Figure S3. Effect of memorandum encoding on distractor-induced bias.**

**A.** Same as in Figure 2C (main text), but showing behavioral bias curves for the cued memorandum based on a median split of memorandum encoding strength – quantified with neural decoding accuracy following memorandum presentation (gray shaded bar, Fig. 2B, left) – when the distractor appeared in the same hemifield as the memorandum (n=23). Solid curve and filled violin plot (*inset*): Weaker memorandum encoding trials. Dashed curve and open violin plot (*inset*). Stronger memorandum encoding trials.

**B.** Same as in panel A, but when the distractor appeared in the hemifield opposite to the cued memorandum. Other conventions are the same as in Figures 2C-D (main text).

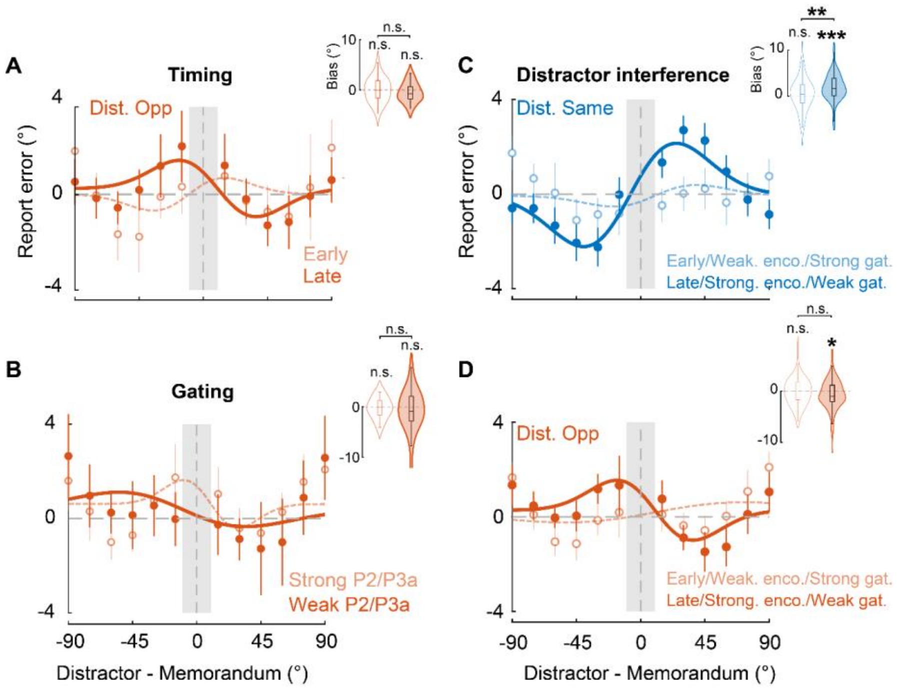


**Supplementary Figure S4. Distractor-opposite and omnibus analysis results.**

**A.** Same as in Figure 4B-C (main text) but showing the behavioral bias curves for the cued memorandum when the distractor appeared on the hemifield opposite to the cued memorandum – for trials in which the distractor appeared early (dashed, open symbols, unfilled violins) or late (solid, filled symbols, filled violins) during the delay period.

**B.** Same as panel A, but for trials with strong (dashed) or weak (solid) distractor-evoked P2/P3a amplitude. (A-B) Other conventions are the same as in Figures 4B-C.

**C.** Same as panel Figure 4B (main text), but showing the results of an omnibus analysis combining across conditions with high distractor interference – late timing, strong encoding, weak gating (solid, filled symbols, filled violins) – or low distractor interference – early timing, weak encoding, strong gating (dashed, open symbols, unfilled violins) – for trials in which the distractor appeared on the same hemifield as the cued memorandum.

**D.** Same as panel C, but when the distractor appeared on hemifield opposite to the cued memorandum. (C-D), other conventions are the same as in Figure 4B (main text) and SI Figure S4A. (All panels) *: $p<0.05$; **: $p<0.01$; ***: $p<0.001$; n.s.: not significant.

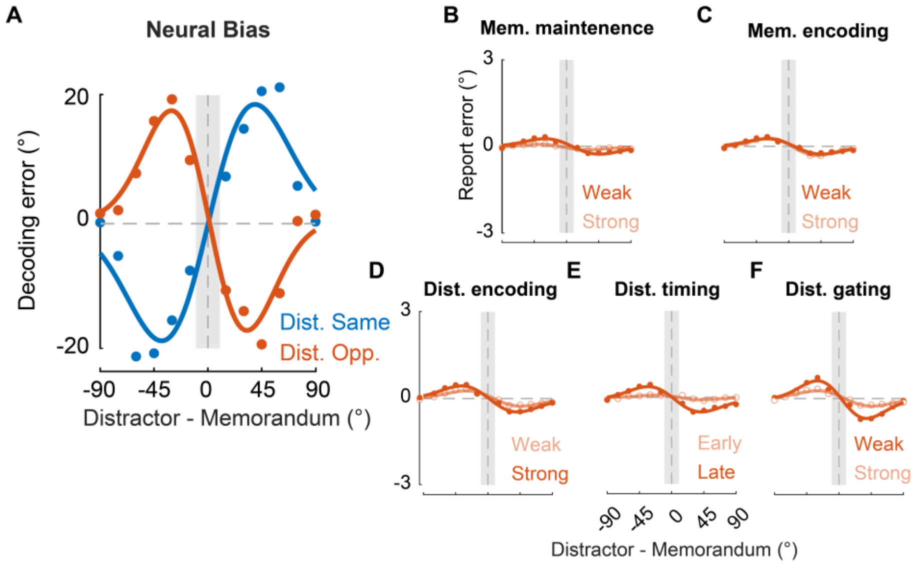


**Supplementary Figure S5. Simulations of neural bias and distractor-opposite effects.**

**A.** Simulated neural bias for the cued memorandum when the distractor appeared in the same hemifield (blue) or the opposite hemifield (orange) as the former. Other conventions are the same as in Figure 6A (main text).

**B-F.** Same corresponding panels as in Figure 6B-F (main text), except showing simulated behavioral bias when the distractor appeared in hemifield opposite to the respective memorandum. Other conventions are the same as in corresponding panels in Figures 6B-F.

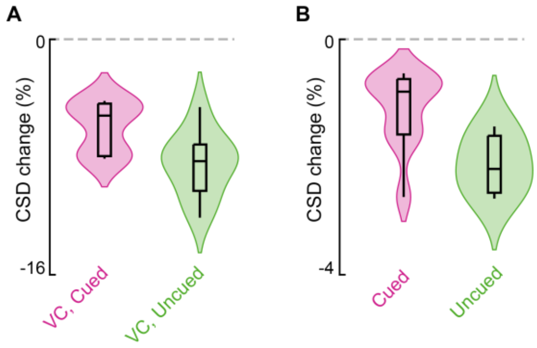


**Supplementary Figure S6. Error-correcting dynamics in WM.**

**A.** Change in error in the VC attractor readout upon incorporating top-down feedback from the HC attractor. Data shown separately for the cued (magenta) and uncued (green) attractors. Violin plots showing the distribution of values across 10 runs of the model (n=1000 simulations for each run); each simulation initialized with different random seeds, and values were averaged across simulations within a run.

**B.** Change in error obtained by averaging readouts from the VC and the HC attractor relative to the mean readout error for either attractor (average of VC and HC readout errors). Other conventions are the same as in panel A.

**Supplementary Table S1. Network architecture and simulation parameters employed in the model.**

**A. Network architecture parameters**

| Parameter | Description | Value |
|---|---|---|
| $W_I^{VC,U \rightarrow VC,C}$ | Inter-hemispheric inhibitory connection strength from uncued visual cortex to cued visual cortex | 0.02 |
| $W_I^{VC,C \rightarrow VC,U}$ | Inter-hemispheric inhibitory connection strength from cued visual cortex to uncued visual cortex | 0.02 |
| $W_E^{HC,C \rightarrow VC,C}$ | Top-down connection strength from higher cortex to visual cortex on the cued hemifield before/after cue onset | 0.1/0.12 |
| $W_E^{HC,U \rightarrow VC,U}$ | Top-down connection strength from higher cortex to visual cortex on the uncued hemifield before/after cue onset | 0.1/0.1 |
| $W_E^{VC,C \rightarrow HC,C}$ | Bottom-up connection strength from visual cortex to higher cortex on the cued hemifield during the stimulus/delay periods | 3.0/0.0 |
| $W_E^{VC,U \rightarrow HC,U}$ | Bottom-up connection strength from visual cortex to higher cortex on the uncued hemifield during the stimulus/delay periods | 3.0/0.0 |
| $W_{EE}$ | Strength of local recurrent connections among excitatory neurons | 6* |
| $W_{EI}$ | Strength of global recurrent connections from excitatory to inhibitory neurons | 4* |
| $W_{IE}$ | Strength of global recurrent connections from inhibitory to excitatory neurons | 3.4* |
| $W_{IE}$ | Strength of global recurrent connections among inhibitory neurons | 0.85* |

*these parameter values were identical for all four attractors (VC,C; VC,U; HC,C; HC,U)

**B. Simulation parameters**

| Parameter | Description | Value |
|---|---|---|
| $\tau_E$ | Time constant of E neurons, in ms | 20 |
| $\tau_I$ | Time constant of I neurons, in ms | 10 |
| $A_M$ | Peak amplitude of the von Mises input current profile for the memory array | 200 |
| $A_D$ | Peak amplitude of the von Mises input current profile for the distractor | $10^3$ |
| $\kappa_M$ | Concentration parameter of the von Mises input current profile for the stimulus | 2.7 |
| $\kappa_D$ | Concentration parameter of the von Mises input current profile for the distractor | 2.25 |
| $C_E$ | Constant bias current input to excitatory neurons | 0.2* |
| $C_I$ | Constant bias current input to inhibitory neurons | 0.5* |
| $\sigma_E$ | Standard deviation of input noise for excitatory neurons | 3* |
| $\sigma_I$ | Standard deviation of input noise for inhibitory neurons | 5* |

*these parameter values were identical for all four attractors (VC,C; VC,U; HC,C; HC,U)
